# Metformin increases tauroursodeoxycholic acid levels to improve insulin resistance in diet-induced obese mice

**DOI:** 10.1101/2020.05.26.116715

**Authors:** Ya Zhang, Yang Cheng, Jian Liu, Dan He, Jihui Zuo, Liping Yan, Ronald W. Thring, Mingjiang Wu, Yitian Gao, Haibin Tong

## Abstract

Metformin is widely used to surmount insulin resistance (IR) and type 2 diabetes. Evidence indicates that metformin improves insulin resistance associated with gut microbiota, but the underlying mechanism remains unclear. In the present study, metformin effectively improved insulin sensitivity and alleviated liver inflammation and oxidative stress in high-fat diet (HFD)-fed mice. Metabolomics analysis showed that metformin increased tauroursodeoxycholic acid (TUDCA) levels both in intestinal content and liver by reducing the production and activity of bile salt hydrolase (BSH). We further found that TUDCA was able to antagonize with KEAP1 to prevent its binding to Nrf2 and activate Nrf2/ARE pathway, thereby reducing intracellular ROS and improving insulin signaling. Moreover, metformin increased the proportion of *Akkermanisia muciniphlia* in the HFD-fed mice, while *in vitro* growth curve test confirmed that it’s TUDCA, not metformin, promoted the proliferation of *A. muciniphlia*. Subsequently, TUDCA administration could effectively ameliorate insulin resistance, activate hepatic Nrf2/ARE pathways, and increase the abundance of intestinal *A. muciniphlia* in *ob/ob* mice. These findings reveal that metformin remodels the gut microbiota, reduces oxidative stress and enhances insulin sensitivity partly due to increasing the production of TUDCA. This provides a novel mechanism by which metformin alleviates diet-induced insulin resistance and improves metabolism.

## Introduction

The prevalence of obesity and overweight have been on the rise worldwide^1^. Current predictions indicate that more than 1 billion people will be obese by 2030^2^. Visceral obesity precedes the development of metabolic complications and chronic diseases such as type 2 diabetes mellitus (T2DM)^3^, hepatic steatosis^4^ and cardiovascular diseases^5^. Metformin has been the first-line anti-diabetic drug for more than 60 years because of its excellent hypoglycemic effect and safety profile^1^. Although several studies have explored the mechanisms by which metformin maintains glucose homeostasis, findings from such studies are inconsistent. The prevailing view is that metformin activates AMP-activated protein kinase (AMPK) in the liver, similar to the principle of caloric restriction that relieves symptoms of diabetes^6^. However, metformin is an orally administered drug that reaches high concentrations in the intestine with much lower serum concentrations^7^, thus, the possibility that its metabolic therapeutic mechanism might be due in part to actions in the intestine cannot be ignored. Accumulated evidence shows that a high-fat diet (HFD) reduces the abundance of *Lactobacilli* in the small intestine. However, this can be counteracted by metformin, which increases the proportion of probiotics, thus, relieving the symptoms of HFD-induced diabetes^8^. Although metformin may play a hypoglycemic effect by improving the gut microbiota, oral administration of antibiotics to diabetic mice impairs the hypoglycemic ability of metformin^9^. In addition, while metformin can increase the abundance of *Escherichia coli*, it can as well reduce the abundance of *Intestinibacter*^10^. This can further influence diverse biological processes such as the synthesis of branched-chain amino acid and the secretion of GLP-1, thereby improving glucose tolerance and alleviating insulin resistance (IR)^10, 11^. These findings suggest that modulation of gut microbiota is involved in the hypoglycemic mechanism of metformin.

Bile acids (BAs), which are produced in the liver and serve as important ingredients of the digestive fluid in the intestine, enter the small intestine through the biliary system and participate in fat digestion and absorption^12^. Gathered evidence demonstrates that BAs could improve the intestinal homeostasis by modulating the gut microbiota, thus relieving metabolic syndrome^13^. Recent studies also suggest that BAs can participate in the liver-gut axis circulation and the improvement of HFD-initiated metabolic disorders by inhibiting the proliferation of pathogenic bacteria^14^. By modulating the gut microbiota, BAs have also been reported to induce multiple beneficial effects on the intestinal lumen and intestinal walls such as the suppression of intestinal inflammatory responses^15^, the resolution of endoplasmic reticulum (ER) stress in the intestinal epithelial cells underlying the pathology of inflammatory bowel disease^16^ and the improvement of gut barrier dysfunction^17–19^. These findings suggest that there is a complex relationship among BAs, gut microbiota and metabolic syndrome.

Previous studies have reported the association of BAs with oxidative stress, which has been shown to treat metabolic dysfunction by acting as an endogenous chemical chaperone to protect cells against ER stress^20^. Intestinal micro-environment (including inflammation status, the function of the epithelial tight junction and gut microbiota) plays an essential role in the progression of obese-induced IR^21^. The presence of gut microbiota and BAs in the intestine may be closely associated with intestinal metabolic state^22, 23^. Analyses from metagenomic sequencing and metabolomics show that metformin treatment increases the levels of the conjugated bile acid in the gut by decreasing the abundance and bile salt hydrolase (BSH) activity of *Bifidobacterium* species in the intestines of individuals with T2DM^24, 25^. BAs are signal molecules that control the dynamic balance between energy metabolism and liver protection. It has been reported that BAs change with obesity, IR^26^ and nonalcoholic steatohepatitis (NASH)^27^. However, the mechanism by which BAs alleviate HFD-induced IR and the underlying role played by the gut microbiota during this process are not clear.

Accumulating evidence has clearly indicated that oxidative stress plays a major role in the pathological process of IR^28^. What’s more, mitochondrial dysfunction, reactive oxygen species (ROS) over production, and lipid peroxidation have been found in the liver of Zucker rats with T2DM^29^. Increased oxidative stress seems to be a deleterious factor leading to IR and impaired glucose tolerance in T2DM^30^. Nuclear factor-erythroid-2-related factor 2 (Nrf2), tightly interacts with Kelch-like ECH-associated protein 1 (KEAP1), an important transcription factor responsible for inducing phase II detoxifying and antioxidant enzymes, is a key player in the antioxidant response and glucose metabolism^31, 32^. Recent study has confirmed KEAP1-Nrf2 system as a critical target for preventing the onset of IR^33^, and bile acids have a great influence on the Nrf2/ARE antioxidant signal pathway^34^. This study investigated how metformin prevents the development of IR in HFD-induced obese mice by modulating the dysbiosis of gut microbiota and alteration metabolites. Exploring the mechanism behind this improves our knowledge of gut microbiota and metabolic interactions underlying the anti-insulin resistance effects of metformin.

## Results

### 1. Metformin alleviates HFD-induced insulin resistance and oxidative stress

The administration of metformin significantly alleviated diet-induced increase of body weight (**Fig. 1A**), without affecting food intake (**Fig. S1A**). The reduced liver index and subcutaneous fat index (**Fig. S1B)** as well as H&E staining of subcutaneous adipose tissue (**Fig. S1C)** and number of adipocytes (**Fig. S1D)** suggested that metformin prevented fat deposition in HFD-fed mice. The diet-induced dyslipidemia was effectively prevented by metformin as indicated by the lower levels of T-CHO, TG and FFA in both the liver and the serum (**Fig. S1E-G**). Further, metformin significantly improved HFD-feeding impaired glucose tolerance and insulin resistance determined by IPGTT (**Fig. 1B**) and IPITT (**Fig. 1C**), respectively. Marked elevation in fasting serum insulin (FINS) was observed in the HFD group. Notably, metformin treatment tended to reverse the increase in FINS (**Fig. 1D**) as well as reduce HOMA-IR (**Fig. 1E**), an indicator for evaluating the level of insulin resistance. In addition, the level of phosphorylated Akt at Ser473 was significantly reduced in the liver of the HFD-induced obese mice, but the level was restored in the HFD+Met group (**Fig. 1F**). Contrary to the phosphorylated Akt, the level of phosphorylated IRS-1 at Ser307 was significantly elevated in the HFD group, whereas this level was reduced in the HFD+Met group (**Fig. 1F**). These results indicate that the preventive effect of metformin on HFD-induced obesity resulted in improved glucose homeostasis and insulin sensitivity.

**Figure 1.**
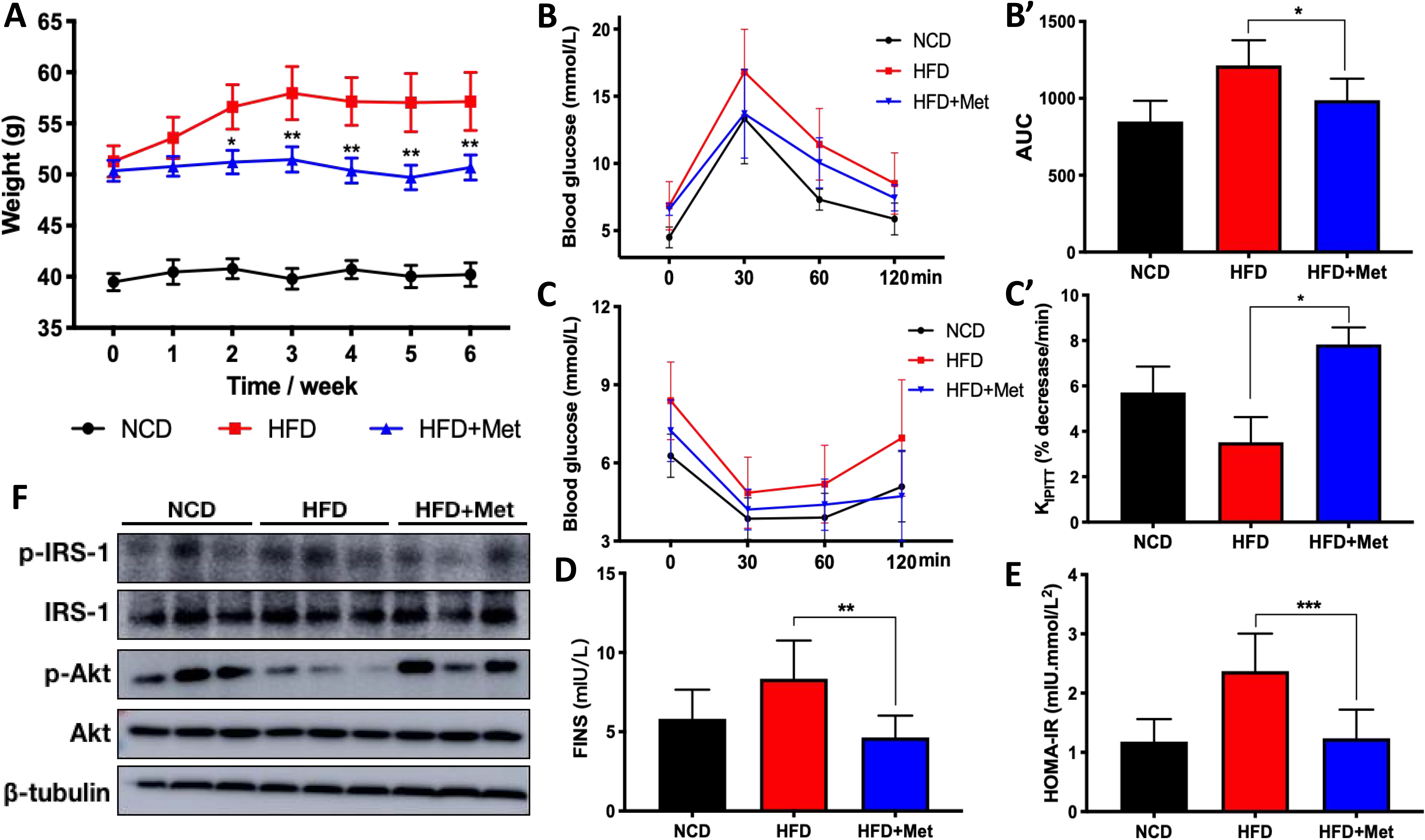
Metformin prevents obesity and insulin resistance in HFD-fed mice. (A) Body weight was shown throughout 6 weeks. (B,C) Mice, fasted overnight and fasted for 6 hours, were injected with glucose (2 g/kg in saline) or insulin (0.75 IU/kg in saline) for IPGTT and IPITT, respectively. (B) IPGTT and (B’) AUC of IPGTT; (C) IPITT and (C’) slope of IPITT. (D) FINS and (E) HOMA-IR. (F) Protein expression of Akt, phosphorylated Akt at Ser473, IRS-1, phosphorylated IRS-1 at Ser307 and quantitative analysis for the densitometry. Data are expressed as mean ± SD. \**P* < 0.05, \*\**P* < 0.01 and \*\*\**P* < 0.001 for HFD *vs* HFD+Met. AUC, area under the curve; FINS, fasting serum insulin; HOMA-IR, homeostatic model assessment for insulin resistance; IPGTT, intraperitoneal glucose tolerance test; IPITT, intraperitoneal insulin tolerance test; Met, metformin.

Accumulated evidence indicated that oxidative stress is involved in the development of HFD- induced IR, thus we further measured the oxidative stress levels after metformin treatment. The results showed that the activity of CAT in the serum and the liver was markedly lower in the HFD group compared with the NCD group (**Fig. 2A**), the ratio of GSH/GSSG was also reduced (**Fig. 2B**). After metformin intervention, the activity of CAT and the ratio of GSH/GSSG were significantly increased. The accumulation of lipid peroxide MDA in liver, caused by HFD, was significantly reduced in the metformin group (**Fig. 2C**). Nrf2/ARE is a classic antioxidant-signaling pathway and KEAP1 is a key protein that mediates the degradation of Nrf2. HFD feeding significantly inhibited Nrf2/ARE signaling whereas metformin significantly increased the expression of Nrf2 as well as phosphorylated Nrf2 (**Fig. 2D**). Nrf2/ARE signaling regulates the expression of a variety of detoxification and antioxidant enzymes. As shown in **Fig. 2E**, Metformin significantly up-regulated the mRNA levels of *Nqo1* and *Ho1*, which are essential genes for balancing oxidative stress. Also, metformin prevented metabolic inflammation in the liver and ileum, leading to a reduced gene expression of *Il-1β*, *Il-6* and *Tnf-α* in the ileum (**Fig. S2A,B**). Metformin also prevented the release of endotoxin LPS in serum (**Fig. S2C**). Meanwhile, the phosphorylated levels of p65 and IκB were decreased in the livers of metformin-treated mice compared with the HFD-fed mice (**Fig. S2D**), suggesting that the HFD-induced activation of NF-κB signaling was suppressed by metformin. These results indicate that metformin can improve HFD-induced insulin resistance, oxidative stress and inflammation.

**Figure 2.**
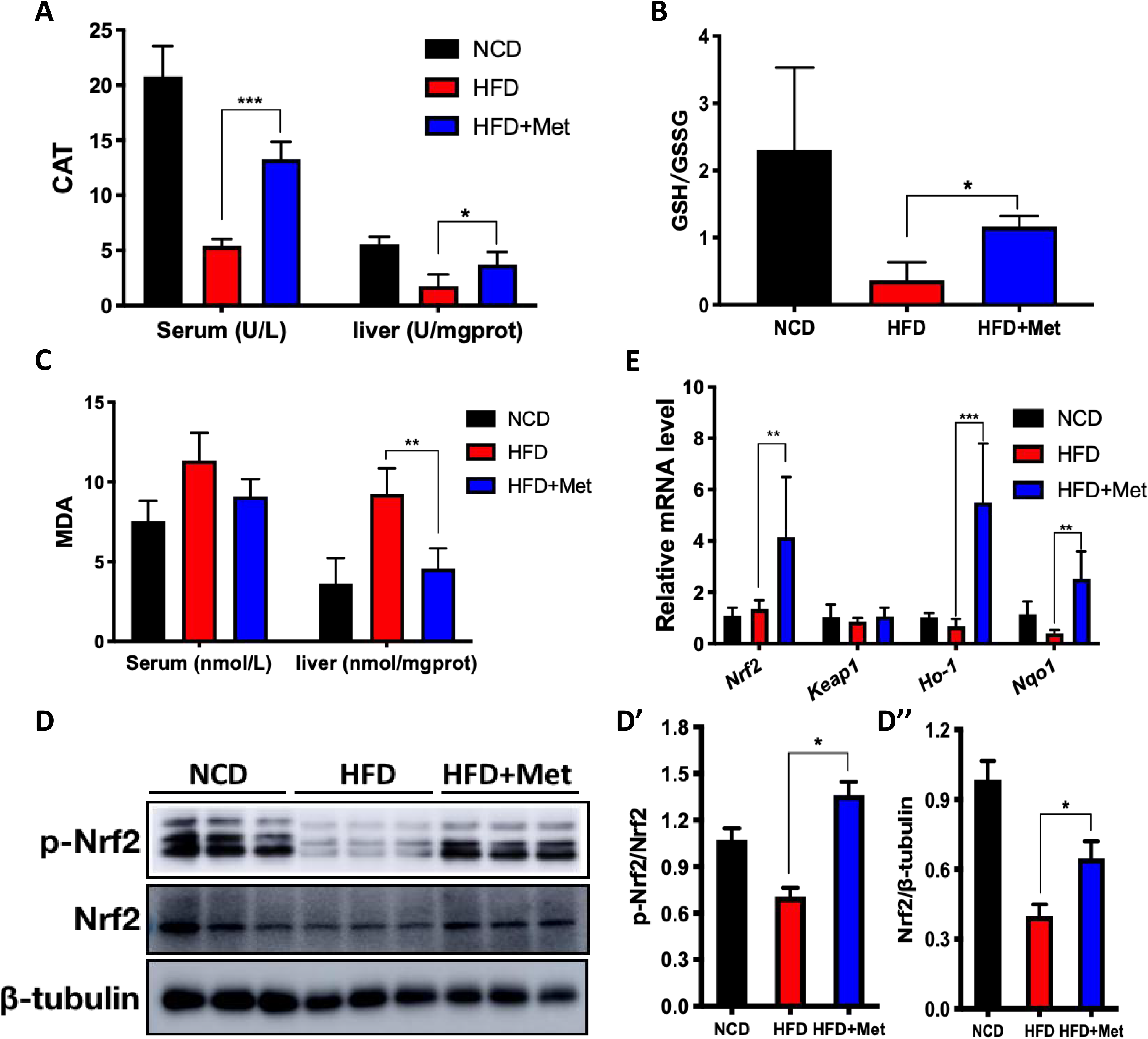
Metformin ameliorates HFD-induced oxidative stress. (A-C) Serum or liver homogenate were used to quantify the representative of oxidative stress: (A) CAT, (B) GSH:GSSG ratio, and (C) MDA. (D) Protein expression of total Nrf2, phosphorylated Nrf2 and (D’, D’’) quantitative analysis for the densitometry. (E) Quantitative RT-PCR analysis of *Nrf2*, *Keap1*, *Nqo1* and *Ho-1* are shown. Data are expressed as mean ± SD. \**P* < 0.05, \*\**P* < 0.01 and \*\*\**P* < 0.001 for HFD *vs* HFD+Met. CAT, catalase; GSH, glutathione; GSSG, oxidized glutathione; MDA, malondialdehyde.

### 2. Metformin prevents obesity-driven dysbiosis of gut microbiota and reshapes intestinal metabolites

Faecal samples were collected for gut microbiota analysis by 16S rRNA amplicon sequencing. Following all quality trimming and checking, three dataset with highly qualified reads were collected for subsequent analysis. Our data showed that the numbers of unique OTUs in the NCD, HFD, and HFD+Met groups were 372, 350 and 374, respectively. The coverage of sequencing results was nearly complete for all sequences in the three groups, demonstrating sufficient sequencing depth for follow-up analyses. The metformin-treated group significantly increased the Ace and Chao1 indices compared with the HFD group (**Table 1**). Metformin-treated mice, however, clustered partially apart from HFD-fed mice samples, suggesting important changes in gut microbial profile (**Fig. 3A, Fig. S3A**). Metformin treatment prevented HFD-induced decrease in Bacteroidetes and the increase in the Firmicutes/Bacteroidetes ratio (**Fig. S3B,C**), two hallmarks of obesity-driven dysbiosis. Based on FunGuild and FAPROTAX analysis of flora function, the HFD group showed a decrease in the proportion of the chemoheterotrophy and the fermentation bacteria when compared with the NCD group (**Fig. S3D**). However, metformin supplementation reshaped the structure of gut microbiota, chemoheterotrophy and fermentation improved in the HFD+Met group. Heatmap presents the clustering of bacterial communities with their relative abundances at the genus level (**Fig. 3B**). Compared with the NCD group, unique high-abundance bacteria in the HFD group are mainly concentrated in *Bifidobacterium, Roseburia, Lachnoclostridum, Alloprevotella* and *Mucispirilum*. However, metformin decreased the abundance of these bacteria, making the metformin-treated group comparable to the NCD group. In addition, metformin specifically increased the abundance of *Butyricimonas, Parasulterella, Parabacteroides* and *Akkermansia*. Consistent with the decrease in Firmicutes/Bacteroidetes ratio, the difference between the HFD group and the HFD+Met group is mainly reflected in the decrease in the abundance of Firmicutes and the increase in the abundance of Bacteroidetes (**Fig. S3E**). Compared with the NCD and HFD+Met groups, *c_Erysipelotrichia, f_proteobacteria, g_Turicibacter, o_Bifidobacteriales* and *s_Clostridium* in the HFD group were significantly up-regulated, whereas f*_Muribaculaceae, g_parabacteroides* and *f_tannerellaceae* were significantly up-regulated in the HFD+Met group. Therefore, metformin can increase the abundance of Bacteriodia at the phylum level and *Akkermansia* at the genus level, and modulate HFD-induced dysbiosis of gut microbiota.

**Figure 3.**
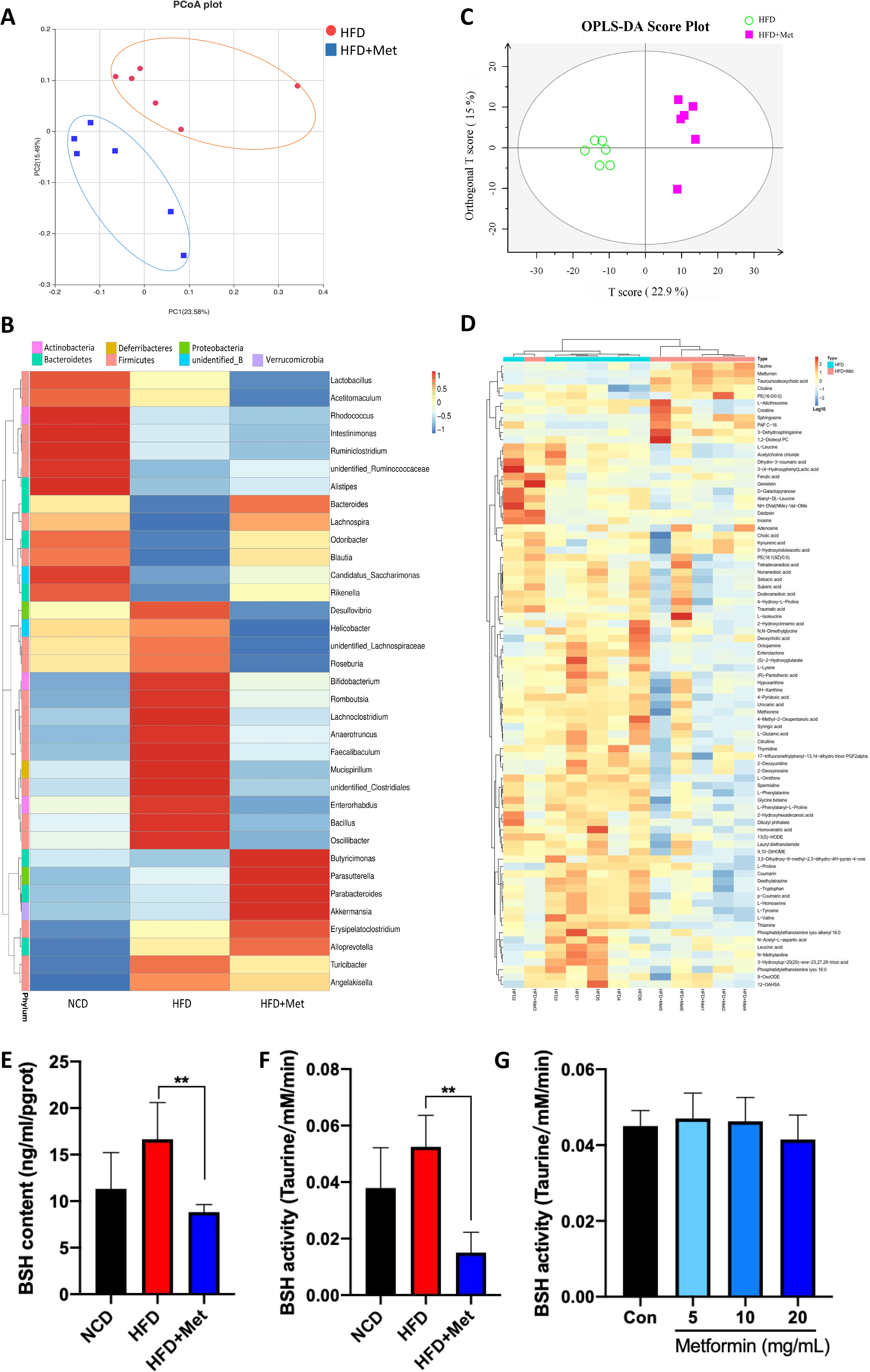
Genus-based comparison of gut microbiota and non-targeted metabolome analysis. (A) Principal coordinates analysis. (B) Heatmap showing clustering of bacterial communities with their relative abundances at the genus level, the bacterial communities or OTUs were clustered phylogenetically by neighbor-joining method and the libraries were clustered by profile pattern using Unifrac analysis. Relative abundances (log values) of microbial genus are displayed in a red-to-blue color code (high to low abundance). (C) OPLS-DA score plot of non-targeted metabolome. (D) Heatmap of 85 significantly differential metabolites. (E) BSH content and (F) BSH activity in faeces of the three groups of mice. (G) Fecal suspensions were prepared by collecting fresh faeces, adding metformin to final concentrations of 5, 10 and 20 mg/mL, incubating for 30 min and then measuring BSH activity. Data are expressed as mean ± SD. \**P* < 0.05, \*\**P* < 0.01 and \*\*\**P* < 0.001 for HFD *vs* HFD+Met. PCoA, principal coordinates analysis; OPLS-DA, orthogonal partial least-squared discriminant analysis; Met, metformin; BSH, bile salt hydrolase.

**Table 1.**
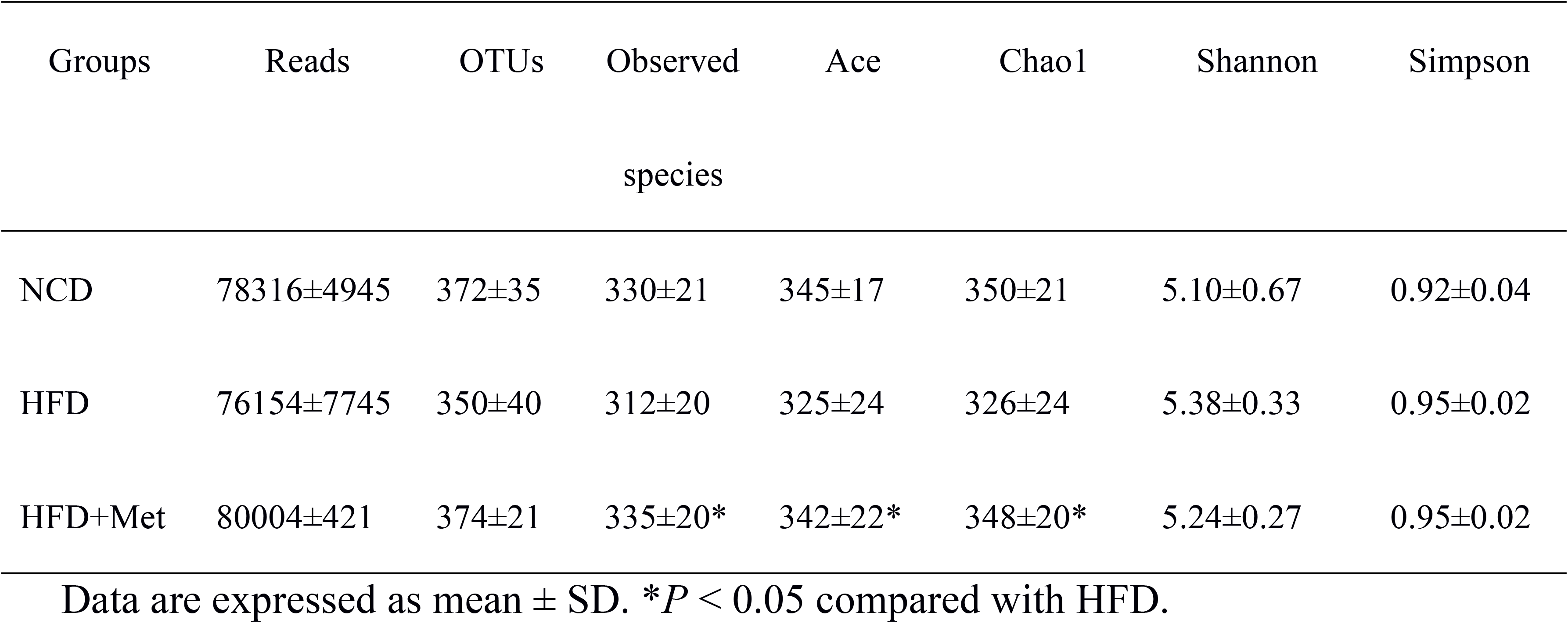
Diversity of gut microbiota in HFD and HFD+Met treated mice

The intestinal metabolite profiles were further measured using lipid chromatography and mass spectrometry (LC-MS) to reveal the influences of metformin on the intestinal metabolites. Metformin caused differential changes of intestinal metabolites in HFD-fed mice evidenced by the scatter plots of OPLS-DA (**Fig. 3C**). Compared to HFD group, metformin up-regulated 182 of intestinal metabolites and down-regulated 118 metabolites in the HFD+Met group (**Fig. S4A**). A total of 85 significantly differential metabolites between HFD and HFD+Met groups were shown in **Fig. 3D**. The HFD+Met group effectively increases the content of taurine, choline and TUDCA in the intestine compared with the HFD group (**Fig. S4B)**. To further determine the source of TUDCA, the TUDCA content both in serum and liver was detected. We found that TUDCA in the serum and liver of the HFD+Met group was significantly increased compared with that in the HFD group (**Fig. S4C)**. In the HFD+Met group, the gene expression of *Cyp7a1* and *Baat*, the important enzymes involved in the biosynthesis of TUDCA, was significantly up-regulated compared with that in the HFD group (**Fig. S4D**). In addition, BSH secreted by *Bifidobacterium* and *Lactobacillus* could degrade the conjugated bile acids, such as TUDCA. The relative abundance of these two bacteria was significantly reduced in the HFD+SFF group compared to that in the HFD group, with a concomitant reduction both in content (**Fig. 3E**) and activity (**Fig. 3F**) of BSH. It is noteworthy that the reduced BSH activity was not caused by metformin (**Fig. 3G**).

**Fig. 4A**,**B** showed the correlation between intestinal metabolism and the gut microbiome at the phylum and species level, respectively, after the intervention of metformin in HFD-fed mice. TUDCA was highly associated with *A. muciniphila* species belonging to the *Verrucomicrobia* phylum. Interestingly, through the analysis of growth curves, it was not metformin but TUDCA significantly accelerated the proliferation of *A. muciniphila in vitro* (**Fig. 4C**). These data suggest that metformin may indirectly improve insulin resistance caused by a high-fat diet through modulation of the intestinal abundance of *A. muciniphila* and content of TUDCA.

**Figure 4.**
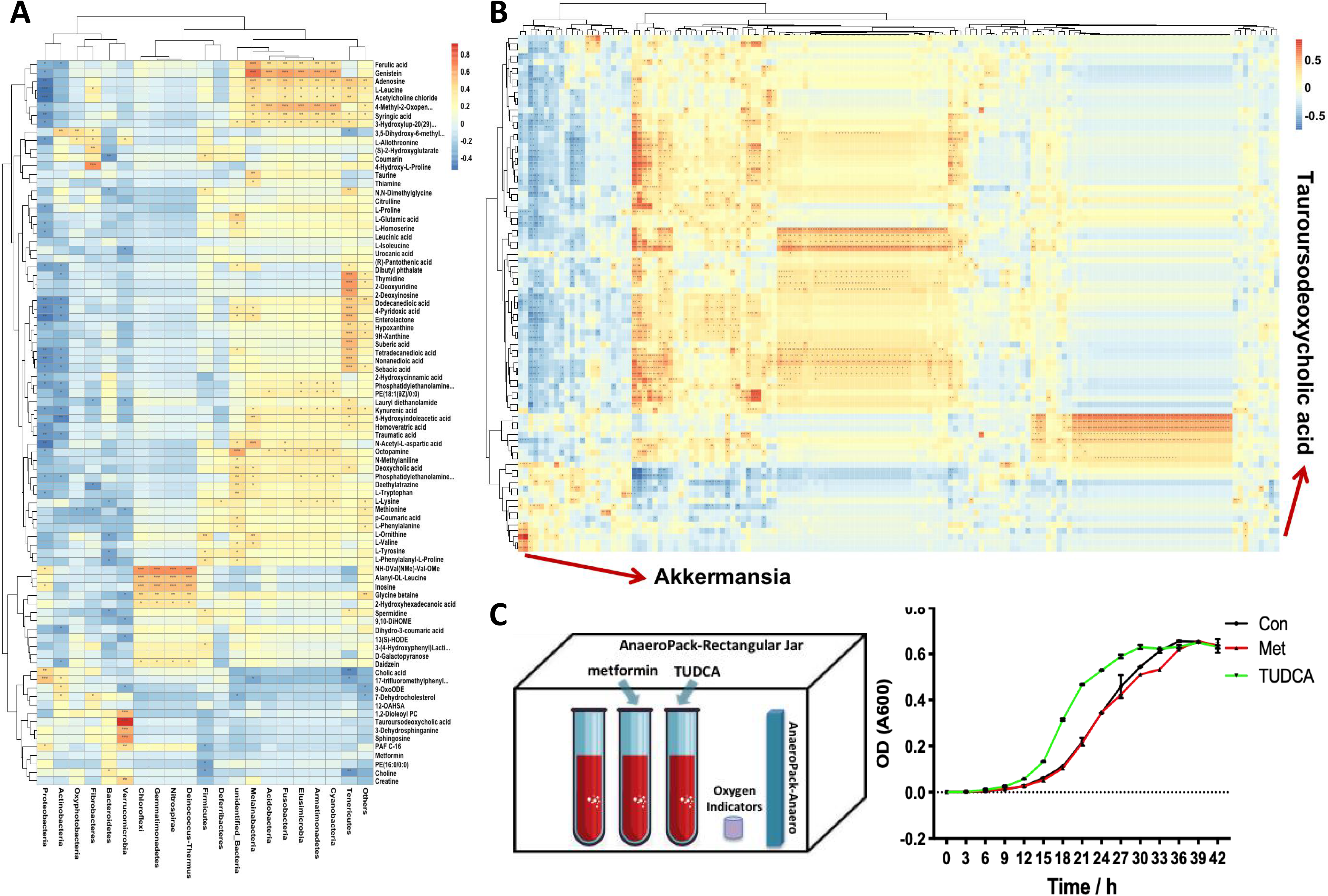
Correlation heatmap for gut microorganisms and metabolites. (A-B) The correlation coefficients of the differential metabolites and gut microbiota were calculated by Pearson’s algorithm using R language, and heat maps were created based on the correlation coefficients. Matrix heatmap showing the correlation between gut microbiota and metabolome at (A) Phylum and (B) Species levels, respectively. (C) Growth curve of *A. muciniphila* in the presence of metformin or TUDCA. Met, metformin; TUDCA, tauroursodeoxycholic acid.

### 3. TUDCA improves palmitic acid (PA)-induced insulin resistance and lipid peroxidation

Further study was carried out to determine whether TUDCA could alleviate hepatic insulin resistance using a PA-treated HepG2 cell model. PA exposure (200 μM) resulted in a decrease in glucose uptake of HepG2 cells, demonstrated by the reduction of fluorescence density in the cytoplasm (**Fig. 5A**) and the high concentration of glucose in the medium (**Fig. 5B**). In addition, PA decreased the level of phosphorylated Akt at Ser473 and increased phosphorylated IRS-1 at Ser307 (**Fig. 5C**), thus, revealing a typical symptoms of insulin resistance. As shown in **Fig. 5A,B**, the impaired glucose uptake caused by PA was significantly alleviated by TUDCA, excluding choline and taurine (**Fig. S5A-C**). Moreover, the suppressed phosphorylation level of Akt in PA-treated HepG2 cells was elevated by TUDCA, while the phosphorylated IRS-1 at Ser307 was also significantly reduced (**Fig. 5C**). Consistent with the above results, in primary mouse hepatocytes, the PA-imparied glucose uptake was also effectively alleviated and the phosphorylated Akt was increased by TUDCA (**Fig. 5D-F**). These findings confirmed that TUDCA could improve hepatic glucose intolerance and enhance insulin sensitivity of PA-treated hepatocyte.

**Figure 5.**
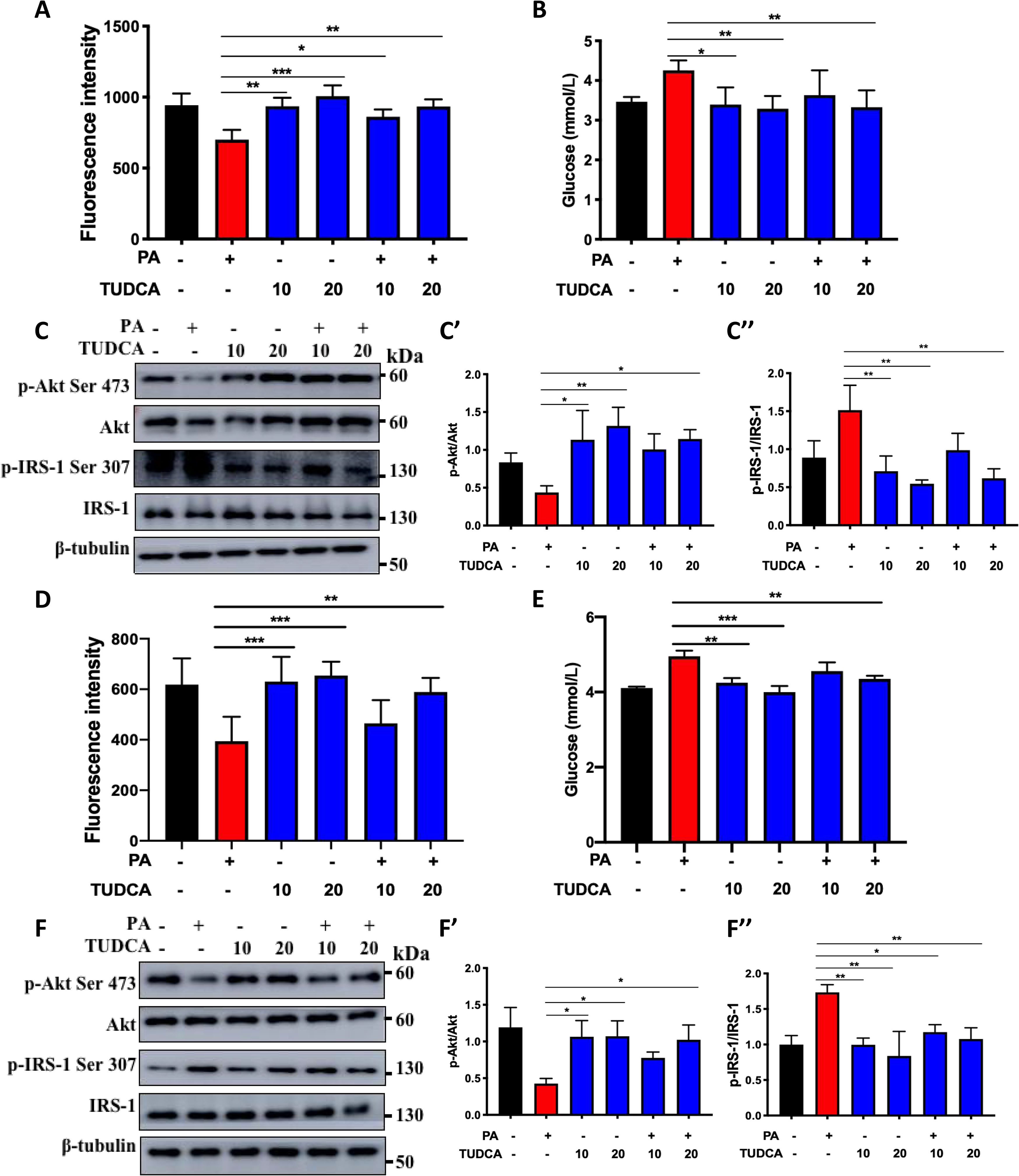
TUDCA alleviates insulin resistance in PA-treated hepatocytes. (A-C) HepG2 cells were pre-treated with 200 μM PA for 24 h and then incubated with TUDCA (10 or 20 μM) for 12 h. (A) Glucose uptake and (B) glucose in the medium supernatant. (C) Protein levels of p-Akt, Akt, p-IRS-1, IRS-1 and (C’, C’’) quantitative analysis for the densitometry. (D-F) Primary mouse hepatocytes were pre-treated with 200 μM PA for 24 h and then incubated with TUDCA (10 or 20 μM) for 4 h. (D) Glucose uptake and (E) glucose in the medium supernatant. (F) Protein levels of p-Akt, Akt, p-IRS-1, IRS-1 and (F’, F’’) quantitative analysis for the densitometry. Data are expressed as the mean ± SD. \**P* < 0.05, \*\**P* < 0.01 and \*\*\**P* < 0.001. PA, palmitic acid; TUDCA, tauroursodeoxycholic acid.

Homeostasis imbalances on glucose metabolism often cause inflammation and oxidative stress damage in hepatic cells. Thus, we further investigated whether TUDCA could alleviate these pathological processes. As shown in **Fig. 6A**, PA could increase ROS generation in primary mouse hepatocytes and HepG2 cells, while TUDCA significantly decreased PA-induced ROS production. On the other hand, the ratio of GSH/GSSG was significantly increased by TUDCA (**Fig. 6B**). In addition, pro-inflammatory cytokines such as *Tnf-α*, *Il-1β* and *Il-6* were also significantly reduced by TUDCA treatment at mRNA level (**Fig. S6**).

**Figure 6.**
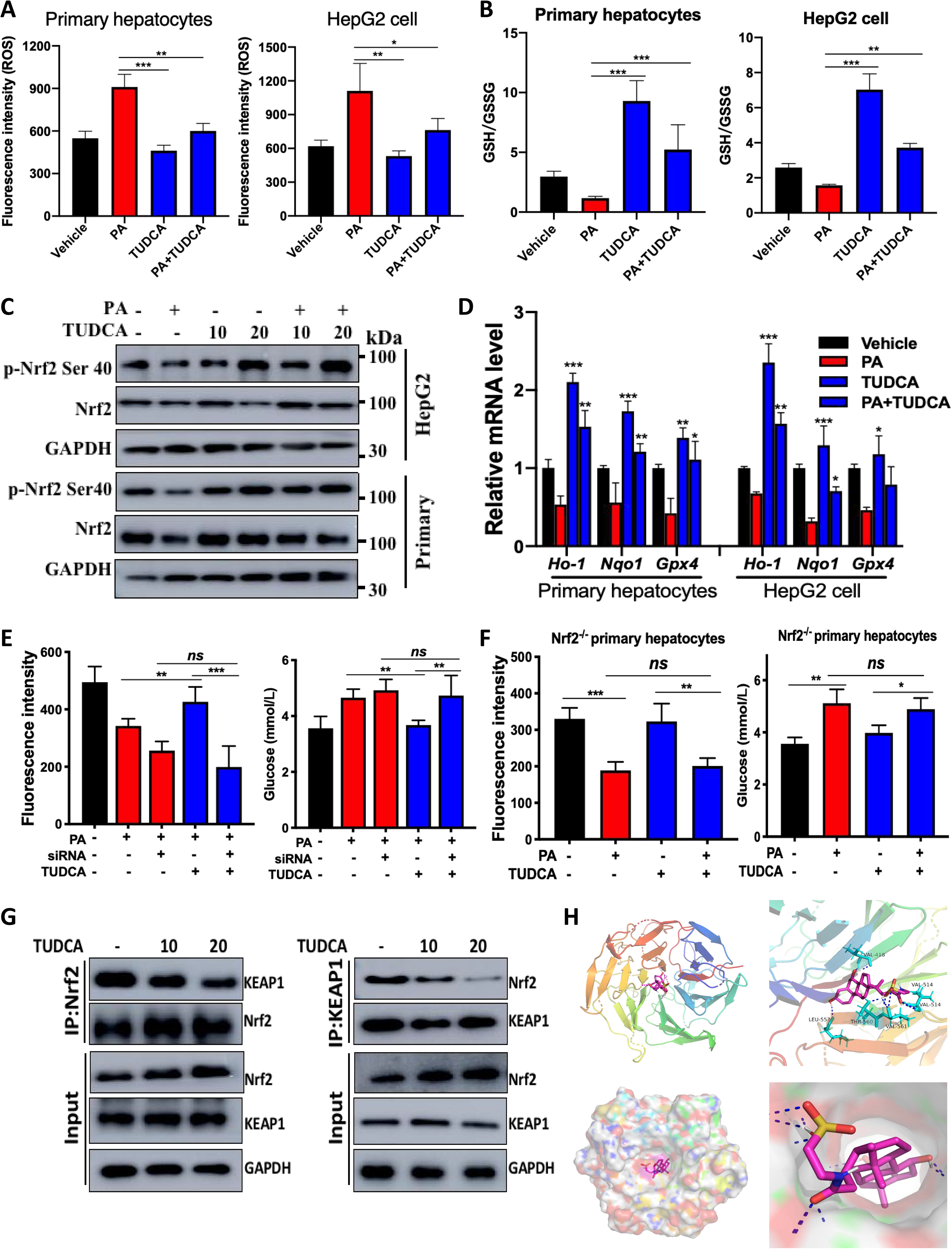
TUDCA relieves PA-induced insulin resistance via activating Nrf2/ARE signaling pathway. (A) Intracellular ROS content; (B) ratio of GSH/GSSG; (C) protein levels of p-Nrf2 and Nrf2 were determined by western blotting in primary mouse hepatocytes and HepG2 cells. (D) Quantitative RT-PCR analysis of *Nrf2*, *Ho-1*, *Nqo1* and *Gpx4* in primary mouse hepatocytes and HepG2 cells. Glucose uptake by cells and glucose content of the medium supernatant measured in Nrf2 knock-down HepG2 cells (E) and primary hepatocytes from Nrf2^-/-^ mice (F). (G) Immunoprecipitation. HepG2 cells were treated with or without TUDCA, and then lysates were immunoprecipitated with the indicated antibodies, followed by western blotting. (H) Docking of TUDCA with Kelch domain of KEAP1. The residues of KEAP1 were represented using sticks or surface structures; TUDCA was shown in pink; the dashed lines (blue) represent hydrogen-bonding interactions. Data are expressed as mean ± SD. \**P* < 0.05, \*\**P* < 0.01 and \*\*\**P* < 0.001. ROS, reactive oxygen species; GSH, glutathione; GSSG, oxidized glutathione; PA, palmitic acid; TUDCA, tauroursodeoxycholic acid.

We further investigated whether TUDCA alleviates PA-induced insulin resistance through the Nrf2/ARE signaling pathway. The transcriptional activity of Nrf2 was determined by measuring the mRNA levels of its representative downstream target genes. TUDCA significantly up-regulated the expression of Nrf2 as well as its phosphorylation level, suppressed by PA (**Fig. 6C, Fig. S7**). It also up-regulated the expression of antioxidant genes, *Ho-1*, *Nqo1*, and *Gpx4*, downstream of the Nrf2/ARE signaling pathway (**Fig. 6D**). We found that TUDCA was unable to alleviate PA-impaired glucose uptake in HepG2 cells that interfered with Nrf2 expression, as well as the primary hepatocytes from Nrf2 knockout mice (**Fig. 6E,F, Fig. S8A,B**). The physical interaction between Nrf2 and KEAP1 was reduced by TUDCA (**Fig. 6G**), suggesting that TUDCA potentially antagonizes with KEAP1 to keep Nrf2 from protein degradation by ubiquitination-proteasome. Binding to the Kelch domain of KEAP1 to achieve the inhibition of KEAP1-Nrf2 interaction is a well-documented mechanism for Nrf2 activation, thus the molecular docking analysis was performed to explore the interaction between TUDCA and the Kelch domain of KEAP1. The theoretical three-dimensional binding mode of the complex with the lowest docking energy is illustrated in **Fig. 6H**. The simulation results revealed that TUDCA was able to interact with the primary amino acid residues on the active site of KEAP1. TUDCA was positioned at the hydrophobic pocket, surrounded by the residues Val-418, Leu-557, Val-514, Val-561, and Thr-560, forming a stable hydrophobic binding. All those interactions helped TUDCA to anchor in the binding site of KEAP1, and the estimated binding energy of KEAP1-TUDCA complex was found to be −10.21 kcal/mol. The docking simulation provided supportive evidence for TUDCA induced Nrf2 activation by allowing us to predict the binding site in Kelch domain.

Further, *ob/ob* mice were employed to validate the effect of TUDCA in alleviating insulin resistance and lipid accumulation *in vivo*. Compared with *ob/ob* mice without treatment, the *ob/ob* mice administrated with TUDCA showed remarkable decrease in body weight, FBG, FINS, as well as HOMA-IR (**Fig. 7A-E**). TUDCA treatment substantially restored glucose intolerance (**Fig. 7F-I**) and insulin resistance (**Fig. 7J**) in *ob/ob* mice. Moreover, similar to WT mice, *ob/ob* mice received TUDCA showed significant amelioration of total cholesterol and triglyceride both in serum and liver (**Fig. S9A,B**). TUDCA significantly improved Nrf2 translocation into the hepatic nucleus (**Fig. 7J**), and up-regulated its downstream antioxidant genes *Nqo1* and *Ho-1* (**Fig. 7K**), suggesting that the Nrf2/ARE signaling pathway was activated by TUDCA. In addition, the abundance of *A. muciniphila* in *ob/ob*+TUDCA group was significantly increased compared to *ob/ob* group (**Fig. 7L**). The gene experssion of pro-inflammatory factors (*Il-1β, Il-6,* and *Tnf-α*) in ileum were also decresed in *ob/ob*+TUDCA group (**Fig. S9C**). Together, these results suggest that TUDCA may significantly activate the Nrf2/ARE signaling, thereby enhancing antioxidant and anti-inflammatory capacity as well as alleviating insulin resistance *in vitro* and *in vivo*.

**Figure 7.**
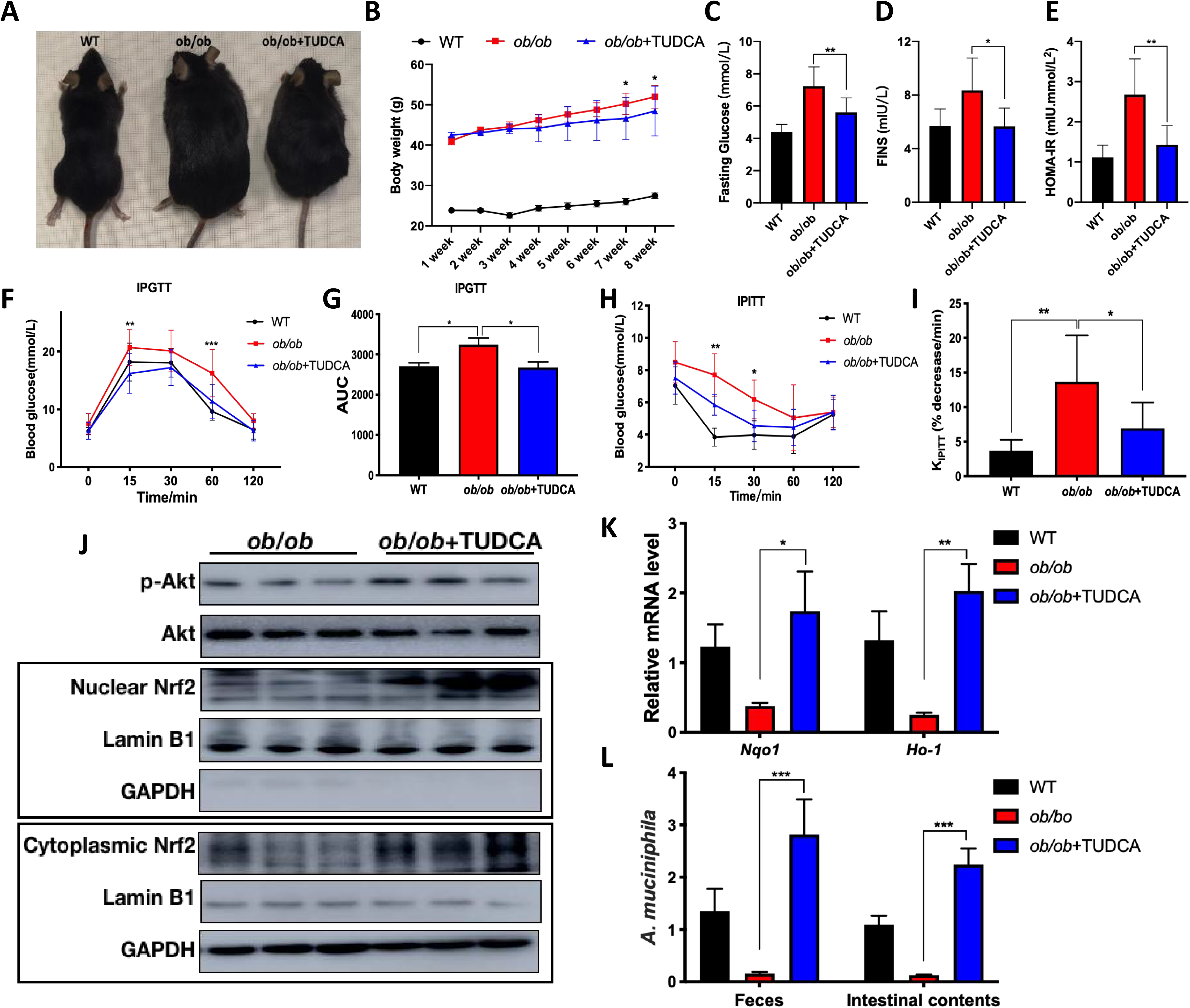
TUDCA improves obesity and insulin resistance in *ob/ob* mice. (A-E) Wild-type C57BL/6J mice and *ob/ob* mice was administered with or without TUDCA for 8 weeks. (A) Body size, (B) body weight, serum levels of (C) FBG and (D) FINS, (E) HOMA-IR. (F-I) Mice, fasted overnight and fasted for 6 hours, were injected with glucose (2 g/kg in saline) or insulin (0.75 IU/kg in saline) for IPGTT and IPITT, respectively. (F) IPGTT, (G) AUC of IPGTT, (H) IPITT and (I) slope of IPITT are shown. (G) Protein expression of p-Akt, Akt, Nrf2 in nuclear and cytoplasm. (K) Quantitative RT-PCR analysis of *Nqo1* and *Ho-1*. (L) Abundance of *A. muciniphila*. Data are expressed as mean ± SD. \**P* < 0.05, \*\**P* < 0.01 and \*\*\**P* < 0.001 for *ob/ob* vs *ob/ob*+TUDCA. AUC, area under the curve; FBG, fasting blood glucose; FINS, fasting serum insulin; HOMA-IR, homeostatic model assessment for insulin resistance; IPGTT, intraperitoneal glucose tolerance test; IPITT, intraperitoneal insulin tolerance test; TUDCA, tauroursodeoxycholic acid.

## Discussion

Metabolic syndromes such as obesity, NAFLD and T2DM are increasingly prevalent globally, especially in recent decades^2^. Metformin, the most widely used anti-diabetic drug worldwide, has been reported to favorably influence metabolic and cellular processes, alleviate insulin resistance^3, 4^; however, the mechanism is still not clear. Recent studies have revealed that the development of metabolic disease strongly correlates to gut microbiota and metabolites. Multiple studies have investigated the impact of metformin on gut microbiota, metabolites and related host targets which regulate glucose metabolism^35^. We explored the potential mechanism involving the gut microbiota and intestinal metabolites to reveal how metformin improves diet-induced obesity and insulin resistance. We established that the daily administration of metformin strongly ameliorated the obesity-related interruption of gut microbiota. The bile acid TUDCA was maintained at a high level in the intestine, which in turn promoted the proliferation of *A. muciniphila*, improved the integrity of the intestinal wall and reduced intestinal inflammation. Furthermore, TUDCA was identified as an activator of Nrf2/ARE signaling in hepatocyte that decreased oxidative stress imbalance, while improving hepatic inflammation and insulin resistance.

There is a variety of microbiota in the human intestine, which is involved in body immunity, infection and nutrient metabolism, and they are closely associated with the occurrence of diabetes. The gut microbiota imbalance in composition and diversity can result in the release of bacterial endotoxin. This could lead to chronic inflammation and destruction of the normal physiological functions of tissues and organs. Several studies have confirmed that chronic inflammation can cause insulin resistance and subsequently diabetes. The decrease in Firmicutes/Bacteroidetes ratio is considered to be an indicator of improved metabolism by metformin therapy^35, 36^. LDA analysis showed that the decreased abundance of Firmicutes by metformin could be due to the decrease in *Erysipelotrichia* species, which belongs to the Firmicutes phylum. The role of *Erysipelotrichia* species in human disease may be well established in studies investigating metabolic disorders^37^. For instance, a previous study showed a bloom of species belonging to the *Erysipelotrichaceae* family (initially classified as *Mollicutes*) in diet-induced obese animals^38^. Within the host, *Erysipelotrichia* species appear to be highly immunogenic^39^, however, in surplus, they induce intestinal inflammation. Studies have shown that species in *Erysipelotrichaceae* are positively correlated with TNFα and IL-6 in the intestine and the liver^40^.

In contrast, most of the bacteria up-regulated by metformin were probiotics. For example, species in *Muribacaceae* are involved in the L-citrulline nitrogen metabolic cycle. They scavenge free radicals, improve immunity and maintain blood glucose. Metformin also up-regulates the abundance of *Parabacteroides*, including *P. distasonis* and *P. goldsteinii*. Liu et al.^41^ found that *P. distasonis* could transform bile acids, activate intestinal gluconeogenesis through succinic acid and improve the host metabolic disorders. In another study, Wu et al.^42^ recorded that *P. goldsteinii* reduces systemic inflammation and increases insulin sensitivity by maintaining intestinal integrity. *Akkermansia muciniphila* is known to effectively alleviate intestinal inflammation caused by obesity^43^. Patients with intestinal obesity have been shown to have a significantly reduced abundance of *A*. *muciniphila* compared with healthy individuals. Numerous studies have demonstrated that metformin could up-regulate the abundance of *A. muciniphila* in HFD-fed mice intestine, maintain the integrity of the intestinal wall as well as effectively improve intestinal inflammation^44^. The abundance of *A. muciniphila* is also significantly up-regulated in the HFD + Met group. As a mucin-degrading bacterium, the use of *A. muciniphila* has been shown to reduce inflammation, maintain the intestinal barrier and increase levels of endocannabinoids secreted by intestinal peptides^45^. We speculate that the transformation of gut microbiota by metformin is the defining factor in improving chronic inflammation in the intestine and even the whole body. Metformin can effectively inhibit the number of pathogenic bacteria and promote the proliferation of probiotics. A suppression of pathogenic bacteria coupled with a proliferation of probiotics may synergistically reduce intestinal inflammation and maintain the integrity and permeability of the intestinal wall. Furthermore, this may lead to a decrease in serum LPS and improvement of systemic inflammation caused by HFD-induced obesity.

The “dialogue” between the gut microbiota and the organism mainly depends on microbial metabolites. By producing various enzymes for biochemical metabolic pathways of the intestinal microbes, gut microbiota can perform diverse metabolic activities such as the metabolism of amino acids, carbohydrates and bile acids, as well as the formation of a co-metabolic relationship with the host. Bile acids produced by the gut microbiota are essential in the regulation of glycolipids and energy. Studies have shown that in human primary hepatocytes, bile acids promote the β-oxidation of fatty acids by activating FXR^46^ and up-regulate the expression of apolipoprotein CII gene to reduce the triglyceride content in the liver^47^.

In our present study, metabolomics data showed that TUDCA in the intestine of HFD+Met group increased significantly. Previous studies revealed that metformin treatment increased the levels of conjugated bile acid in the gut by decreasing the abundance of *Bifidobacterium* and its bile salt hydrolase (BSH) activity in the intestines of individuals with T2DM^14, 25^. BSH, secreted by intestinal bacteria, decomposes TUDCA in the intestine and metabolizes TUDCA to taurine and UDCA. The data from gut microbiota showed that metformin reduced the intestinal bacterial abundance of *Bifidobacterium* and *Lactobacillus* in the mice fed a high-fat diet, while *Bifidobacterium* was the main BSH-producing bacterium. In addition, the content and activity of BSH in the mouse intestine of HFD+Met group were significantly reduced, which was not related to metformin treatment. We hypothesize that high levels of TUDCA are caused by a reduction in BSH-producing bacteria due to the remodeling of the gut microbiota by metformin. *In vitro*, TCDCA can be transformed into TUDCA by 7α/β-hydroxysteroid dehydrogenase (7α/β-HSDH)^12^. Data from the gut microbiota also revealed that metformin did not affect the abundance of *Eubacterium* and *Clostridium*, the major 7α/β-HSDH-producing bacteria. Therefore, the increase of TUDCA in the intestine is purportedly not due to the inhibitory effect of metformin on BSH-producing bacteria. According to the biosynthesis of bile acids in liver, the conjugated bile acid TUDCA can be generated in the liver through a series of catalytic reactions and enzymes such as CYP7A1 and BAAT^12, 18^. CYP7A1 is a rate-limiting enzyme that catalyzes the breakdown of cholesterol into bile acids in the liver^12^, whereas BAAT is a vital enzyme that catalyzes the amidation reaction of taurine and UDCA^48^. Interestingly, we found that the gene expressions of *Cyp7a1* and *Baat* were significantly enhanced by metformin, which might further promote the synthesis of TUDCA in the liver. Thus, we suppose that the increase of TUDCA in the intestine might due to the hepatic biosynthesis by metformin treatment.

TUDCA has been shown to treat NAFLD by acting as an endogenous chemical chaperone; to protect cells against ER stress and to reduce liver steatosis^20^. Moreover, TUDCA can alleviate dextran sulfate sodium-induced colitis in mice^49^. *In vitro*, we found that TUDCA antagonizes with KEAP1 to keep Nrf2 from protein degradation to activate Nrf2/ARE signaling pathway. This is reminiscent of the expression pattern of Nrf2, a crucial stress regulator to reduce ROS, which is a key factor in sensitizing insulin signaling^30, 50^. It has also been confirmed that up-regulation of the Nrf2 expression both by genetic manipulation and pharmacological interference, can significantly decrease the content of ROS and ameliorate IR and T2DM^31–34^. In the mice supplemented with metformin, the significant up-regulation in the expression of *Nrf2* and its downstream target genes (*Gclc*, *Cat* and *Ho-1*) occurred, which further alleviated IR and over-oxidation^51, 52^. Conversely, the stress-resistant effect of TUDCA was dramatically repressed by the inhibition of the Nrf2/ARE signaling. We also found that TUDCA can effectively reduce the inflammation and oxidative stress as well as decreasing the accumulation of lipid peroxide *in vitro* and *in vivo*. This might be a novel mechanism for metformin in the treatment of insulin resistance and diabetes.

Overall, in our present study, metformin was found to increase TUDCA by reducing the production of BSH, which antagonized the interaction between KEAP1 and Nrf2, prevented Nrf2 degradation and promoted its nuclear translocation. TUDCA activated Nrf2/ARE pathway and increased the gene transcription of antioxidant enzyme, further reduced oxidative stress and sensitized insulin signaling. The increase of TUDCA in the intestine corresponded to the increase in the abundance of *A. muciniphlia.* Metformin remodeled the gut microbiota by promoting probiotics such as *Muribacaceae,* and *Parabacteroides*, and inhibiting pathogenic bacteria such as *Erysipelotrichia* and *Helicobacter*, which maintained intestinal integrity and reduced inflammation. However, for a better understanding of the crucial mechanisms involved in the long-term treatment with metformin in humans, the role of the intestine-liver axis should be explored. Since TUDCA is a novel effector molecule of metformin in the gut, plays an important role in alleviating metabolic syndrome.

## Materials and methods

### Chemicals and diet

Metformin (purity ≥ 98%), choline (purity ≥ 98%), taurine (purity ≥ 99%) and palmitic acid (PA, purity ≥ 99%) were obtained from Sigma-Aldrich (St. Louis, MO, USA). Tauroursodeoxycholic acid (TUDCA, purity ≥ 99%) was purchased from Selleck (Houston, Texas, USA). Both the normal chow diet (NCD, containing 10% fat by energy) and high-fat diet (HFD, containing 60% fat by energy) were purchased from Beijing HFK Bio-Technology Co., Ltd. (Beijing, China). Antibodies against Akt, phospho-Akt (Ser473), IRS-1, phospho-IRS-1 (Ser307), KEAP1, GAPDH, lamin B1 and β-tubulin were obtained from Cell Signaling Technology (Beverly, MA, USA). Nrf2, phospho-Nrf2 (Ser40), and 4-HNE were purchased from Abcam (Cambridge, UK). Horseradish peroxidase (HRP)–conjugated secondary antibody was obtained from Santa Cruz Biotechnology (Santa Cruz, USA).

### Animal experiments

All male mice (ICR mice, 7 weeks old, body weight 25 ± 2 g; C57BL/6J mice, 7 weeks old, body weight 20 ± 2 g; *ob/ob* mice, 7 weeks old, body weight 40 ± 2 g; Nrf2^-/-^ mice, 7 weeks old, body weight 20 ± 2 g) were housed in ventilated cages (three animals per cage) at the SPF facility of Wenzhou University under controlled environmental conditions (temperature 22 ± 2°C; relative humidity 60–70%) with free access to standard laboratory chow and tap water. The mice were maintained on a regular 12/12 h light/dark cycle.

Mice were acclimatized to their environment for 1 week before the experiments. Mice were randomly allocated into three groups (n = 9 for each group). A co-worker blinded to the experimental protocol randomized animals into these groups. As for the pharmacological activity of metformin, one group of ICR mice was fed with NCD and the other two groups were fed with HFD. After 12 weeks of HFD feeding, one group of HFD-fed mice was administered metformin (200 mg/kg body weight), once daily (HFD+Met group) for 6 weeks, whereas the NCD-fed mice (NCD group) and other group of HFD-fed mice (HFD group) were treated with an equal volume of saline for 6 weeks. The whole study lasted 18 weeks, during which the body weight, water consumption and food intake were measured every week. At week 18, intraperitoneal glucose tolerance test (IPGTT) and intraperitoneal insulin tolerance test (IPITT) were performed as previously described^1^. As for the pharmacological activity of TUDCA, one group of *ob/ob* mice was administered TUDCA (300 mg/kg body weight), whereas the wild-type C57BL/6J mice and *ob/ob* mice in the other two groups were treated with an equal volume of saline for 8 weeks as control.

Fresh feces were collected and stored immediately at 80°C for subsequent analysis. At the end of the trial, after overnight fasting for 12 h, blood samples were collected, and serum was isolated by centrifugation at 1000 × *g* for 15 min at 4°C, and stored at −80°C for further assay. Tissues, including the adipose tissue, liver and ileum, were weighed; one portion of the tissues was fixed with 10% formaldehyde for histological analysis, and the other portion was immediately frozen in liquid nitrogen for further analysis.

Organ index were calculated by the following formula:

BMI = body weight (kg)/body length (m)^2^

Liver index = wet weight of liver/bw* 100%

Kidney index = wet weight of kidney/bw* 100%

Pancreas index = wet weight of pancreas/bw* 100%

Subcutaneous fat index = wet weight of adipose tissue/bw * 100%

### Biochemical analysis

Total cholesterol (T-CHO), triacylglycerol (TG), free fatty acids (FFA), fasting blood glucose (FBG), fasting serum insulin (FINS), acid phosphatase (ACP), alkaline phosphatase (AKP), alanine aminotransferase (ALT), aspartate aminotransferase (AST), catalase (CAT), malondialdehyde (MDA), lipopolysaccharides (LPS) were determined by biochemical kits purchased from Jiancheng Bioengineering Institute (Nanjing, China). The bile salt hydrolase (BSH) content was determined by biochemical kit purchased from Jianglia Bioengineering Institute (Shanghai, China). The ratio of reduced glutathione (GSH) and oxidized glutathione (GSSG) were determined by a GSH:GSSG kit (Jiancheng Bioengineering Institute, Nanjing, China). Homeostasis model assessment-estimated insulin resistance (HOMA-IR) was calculated using the following formula:

HOMA-IR = FBG (mmol/L) × FINS (mU)/22.5

### Hematoxylin and eosin staining

Adipose tissue were fixed in 10% formaldehyde overnight, paraffin-embedded, sectioned (4-μm thickness, 3–5 sections/specimen) and stained with hematoxylin and eosin (H&E) for histological analysis. Digital images of H&E stained sections were acquired with a Nikon Eclipse Ti microscope at ×400 magnifications (Ti-E/U/S, Japan). Image J software (National Institutes of Health, USA) was used to count adipocytes.

### Cell culture

Human liver hepatocellular carcinoma (HepG2) cells were cultured in MEM (Sigma, St. Louis, MO, USA) supplemented with 10% FBS, penicillin and streptomycin. Cells were maintained in 5% CO_2_ at 37°C.

### Isolation and culture of primary hepatocyte

Male C57/B6J and Nrf2^-/-^ mice (8 weeks old) were used for isolating primary hepatocyte. Hepatocytes were isolated by a two-step collagenase perfusion technique. Briefly, the inferior vena cava was cannulated with angiocatheter and the portal vein was cut. The liver was perfused via the inferior vena cava with 100 mL of PBS at 37 ℃, followed by perfusion with 100 mL of collagenase type IV (Wellington) in HBSS containing Ca^2+^ and Mg^2+^ (GIBCO). After the liver was digested, it was dissected out and cut into small pieces and passed through a 100 μm strainer (Falcon). Hepatocytes were separated from non-parenchymal cells by low-speed centrifugation, and further purified by Percoll gradient centrifugation (50% v/v, Sigma). Cells were plated at a density of 0.3 × 10^6^ on 6-well collagen-coated plate. Hepatocytes were allowed to recover overnight and experiments were started 24 h post isolation.

### siRNA transfection

*Nrf2* was knocked-down by RNA interference (RNAi) using the following 19-bp (including a 2-deoxynucleotide overhang) siRNAs (Origene, Beijing, China): *Nrf2*, SR321100A-AUUGAUGUUUCUGAUCUAUCACUTT; SR321100B-GUCAGUAUGUUGAAUCAGUAGUUTC; SR321100C-CCAGUCUUCAUUGCUACUAAUCAGG. Stealth RNAi (Origene, Beijing, China) was used as a negative control (siCont). For transfection, cells were seeded on a six-well plate, grown to ∼80% confluence and transfected with siRNA duplexes using Lipofectamine 3000 (Invitrogen, Camarillo, CA, USA) according to the manufacturer’s recommendations. After incubation for 48 h, the expression level of Nrf2 protein was detected by western blotting.

### Glucose uptake assay

Glucose uptake assay was carried out according to the previous study with some modifications^53^. Cells were plated at 1 x 10^4^/well in 96-well plates and used at subconfluence after 24 h of preincubation. For experiments, all culture medium was removed from each well and replaced with 100 µM fluorescent 2-Deoxy-2-[(7-nitro-2,1,3-benzoxadiazol-4-yl) amino]-D-glucose (2-NBDG) in serum-free medium and incubated for 30 min. Subsequently, cells were washed thrice with PBS and then the fluorescence intensity was determined with a fluorescence microplate reader (Ex/Em, 488/520 nm). The glucose concentration in the medium supernatant was determined by a glucose detection kit (Jiancheng Bioengineering Institute, Nanjing, China).

### Determination of ROS levels

Intercellular ROS levels were measured using a ROS Assay Kit (Beyotime, Shanghai, China) according to the manufacturer’s protocol. In brief, the dichlorodihydrofluorescein diacetate (DCFH-DA; 10 mM) supplied in the kit was diluted to 10 µM in serum-free medium. Subsequently, cells were pretreated with TUDCA (20 µM) for 2 hour at a density of 3 × 10^3^/96-well. Next, 200 µM PA solution was added and co-incubated for 24 hours. The supernatant was then replaced with serum-free medium. The DCFH-DA solution (10 µM) was added to the cells for 30 minutes at 37°C, and they were then resuspended with PBS (0.1 mM) after washing twice with the indicated inducers. Fluorescence intensity was detected using a hybrid/multi-modereaders (BioTek instruments Inc., Vermont, USA) to determine ROS levels.

### Quantitative RT-PCR

Total mRNA was isolated from tissue samples using TRIzol reagent (TAKARA, Tokyo, Japan) and was reverse-transcribed into cDNA using a high-capacity cDNA reverse transcription kit (TAKARA, Tokyo, Japan) according to the manufacturer’s protocol. The mRNA levels were quantified with quantitative PCR (qPCR) using SYBR Green (Qiagen, Hilden, Germany). Amplification was performed on a LightCycler480 *q*RT-PCR system (Roche, Mannheim, Germany) under the following reactions: 95°C for 15 min, followed by 40 cycles at 95°C for 10 s, 60°C for 20 s, and 72°C for 20 s. The relative mRNA levels of target genes were normalized to the expression of β-actin calculated using 2^−ΔΔct^ method. The primer pairs used in this study are listed in **Supplementary table**.

### Western blotting

Samples were homogenized with ice-cold RIPA lysis buffer containing protease and phosphatase inhibitors (Beyotime, Shanghai, China). The homogenates were centrifuged at 10 000 × *g* for 20 min at 4°C to remove the insoluble tissue debris. The protein concentration in the supernatant was determined using a BCA protein assay kit (Beyotime, Shanghai, China). Equal amount of protein for each group were denatured in boiling water for 5 min. Aliquots (40 μg) of protein samples were subjected to 10% SDS-PAGE and transferred to PVDF membranes (Millipore, Bedford, MA, USA). After blocking with 5% nonfat milk (dissolved in TBST) for 1 h, the membranes were incubated with the indicated antibodies at 4°C overnight, followed by incubation with the appropriate HRP-conjugated second antibodies for 1 h at room temperature. Chemiluminescent detection was performed using the ECL Plus Western blotting reagent (TransStart, Beijing, China). Semi-quantitative analysis for densitometry of each band was performed using ImageJ software.

### Co-immunoprecipitation

For co-immunoprecipitation (Co-IP), the whole protein lysates prepared from HepG2 cells were extracted in a RIPA lysis buffer. Briefly, cell lysates were incubated with Nrf2 or KEAP1 antibodies for 2 h at room temperature, followed by binding of antigen antibody complexes to protein A/G magnetic beads (Thermo scientific, Rockford, USA) for 1 h. Then, the antigen/antibody immunocomplex was stripped by boiling with 5× loading sample buffer for western blotting incubated with KEAP1, Nrf2 and GAPDH antibodies.

### Methodologies of gut microbiota by 16S rRNA amplicon sequencing

Sample preparation: Fæcal samples were freshly collected at week 16 and immediately stored at -80°C. Total fecal DNA was extracted using CTAB/SDS method. The V4 region of the 16S rRNA was amplified using the universal primers 515F and 806R. All PCR reactions were carried out with Phusion^®^ High-Fidelity PCR Master Mix (New England Biolabs), and the mixture PCR products were purified with GeneJET Gel Extraction Kit (Thermo Scientific). Sequencing libraries were generated using Ion Plus Fragment Library Kit (Thermo Scientific) following manufacturer’s recommendations. The library quality was assessed on the Qubit 2.0 Fluorometer (Thermo Scientific). At last, the library was sequenced on an Ion S5 ™XL platform (Thermo Scientific) and 400 bp/600 bp single end reads were generated.

Data acquisition: Paired-end reads were merged using FLASH (V1.2.7), and the Raw fastq files were processed by QIIME (version 1.7.0). Sequence analysis was performed by Uparse software (v7.0.1001), and Sequences with ≥ 97% identity were assigned to the same Operational Taxonomic Units (OTUs).

Data analysis: To compare the compositional OTUs of the gut microbiota in each group, a Venn diagram was constructed using R packages (version 3.1.0) as previously documented. Mothur software packages (version V.1.30.1) were used to calculate the value of Chao1, Ace, Simpson index and Shannon index for evaluation of the community richness and community diversity. Pair-group method with arithmetic means (UPGMA) clustering was generated using the average linkage and conducted by QIIME software (V1.7.0). Linear discriminant analysis (LDA) was carried out to determine the highly dimensional gut microbes and characteristics associated with NCD mice, HFD mice and HFD+Met mice.

### Methodologies of non-targeted metabolome analysis

Metabolites extraction from colon contents: Colon contents were added ddH2O (4°C) and mixed. 100 mg of sample was extracted with 1000 μL of pre-cooled methanol (−20°C). After centrifugation, the supernatant was evaporated and finally dissolved in 400 μL methanol aqueous solution (1:1, 4°C). For the quality control (QC) samples, 20 µL of extract was taken from each sample and mixed. These QC samples were used to monitor deviations of the analytical results from these pool mixtures and compare them to the errors caused by the analytical instrument itself. And the rest of the samples were used for LC-MS detection.

UPLC Conditions: Chromatographic separation was accomplished in an Acquity UPLC system equipped with an ACQUITY UPLC^®^ HSS T3 (150 × 2.1 mm, 1.8 µm, Waters) column maintained at 4°C. The temperature of the auto sampler was 4°C. Gradient elution of analytes was carried out with 0.1% formic acid in water (A) and 0.1% formic acid in acetonitrile (B) at a flow rate of 0.25 mL/min. Injection of 5 μL of each sample was done after equilibration. An increasing linear gradient of solvent B (v/v) was used as follows: 0∼1 min, 2% B; 1∼9.5 min, 2%∼50% B; 9.5∼14 min, 50%∼98% B; 14∼15 min, 98% B; 15∼15.5 min, 98%∼2% B; 15.5∼17 min, 2%.

Mass spectrometry conditions: The ESI-MS^n^ experiments were executed on the Thermo LTQ Orbitrap XL mass spectrometer with the spray voltage of 4.8 kV and -4.5 kV in positive and negative modes, respectively. Sheath gas and auxiliary gas were set at 45 and 15 arbitrary units, respectively. The capillary temperature was 325°C. The voltages of capillary and tube were 35 V and 50 V, -15 V and -50 V in positive and negative modes, respectively. The Orbitrap analyzer scanned over a mass range of m/z 89-1000 for full scan at a mass resolution of 60000. Data dependent acquisition (DDA) MS/MS experiments were performed with CID scan. The normalized collision energy was 30 eV. Dynamic exclusion was implemented with a repeat count of 2, and exclusion duration of 15 s.

Data processing: UPLC-QTOF-MS raw data were analyzed with MarkerLynx Application Manager 4.1 (Waters Corp.). The matrix from UPLC-QTOF-MS was introduced into SIMCA-P 11.0 software (Umetrics) and standardized to a mean of 0 and variance of 1, according to the formula [X - mean(X)]/ std (X), for multivariate statistical analysis. The t test with false discovery rate correction was used to measure the significance of each metabolite. Partial least-squared discriminant analysis (PLS-DA) and orthogonal partial least-squared discriminant analysis (OPLS-DA) were conducted to identify the metabolite discrimination between the two group samples. Differential metabolites were defined with variable importance in the projection (VIP) > 1.0 obtained from OPLS-DA and *P* values less than 0.05 obtained from t test. Differential metabolites were tentatively identified by database matching, i.e., Human Metabolome Database (HMDB) (http://www.hmdb.ca), Metlin (http://metlin.scripps.edu), massbank (http://www.massbank.jp/), LipidMaps (http://www.lipidmaps.org), mzclound (https://www.mzcloud.org). Heatmaps of differential metabolites among all groups were obtained based on spearman correlation and cluster analyses.

### Growth curve of Akkermansia muciniphila

*Akkermansia muciniphila* strain ATCC BAA-835 was purchased from the Beina Biologicals, Inc. (Beijing, China). Bacteria were cultured in BHI medium in tubes at 37°C in an anaerobic chamber (Whitley A35 Workstation 2.5, Don Whitley Scientific, UK). To acquire the growth curve of *A. muciniphila*, the medium in test tubes supplemented with or without 10 mM metformin (or 20 μM TUDCA) was placed in an anaerobic bag to eliminate oxygen for 24 h. The test tubes were then inoculated with bacteria and incubated at 37°C in an anaerobic bag. The OD_600_ of the cultures was measured every 3 hours.

### Determination of hepatic/serum TUDCA

The contents of TUDCA in tissues were determined by high-performance liquid chromatography (HPLC). Briefly, 200 mg of frozen tissue was homogenized with a Qiagen TissueLyserII (Germantown, MD, USA) in 1 mL of PBS to prepare the tissue homogenates. The impurities were removed through a 0.22 μm filter, then the filtrate was precipitated by methanol and 10 µL of supernatant was analyzed using an Agilent 1290 HPLC system (Santa Clara, CA) equipped with a Hypersil ODS-2 column (5 µm, 4.6 × 250 mm; Waters, USA). The supernatant was detected at the wavelength of 210 nm, eluted with the mobile phase of 0.03 M phosphate buffer solution (pH 4.4) and methanol (32:68, v/v) at a flow rate of 1.0 mL/min.

### Determination of bile salt hydrolase activity *invitro*

The enzyme activity of BSH was determined using an amino acid chromogenic method. Briefly, mouse faeces were collected and the faecal suspension was prepared. 10 µL of sample (containing 500 µg of protein) was incubated with 180 µL of sodium phosphate buffer and 10 µL of TDCA (final concentration 10 mM) at 37°C for 30 min. 50 µL of the mixture was mixed with 50 µL of 15% TCA and centrifuged at 10,000g for 10 min. Mix 20 µL of supernatant with 80 µL of water, add 1.9 mL of ninhydrin ( 0.5 mL 1% ninhydrin, 1.2 mL glycerol, 0.2 mL 0.5 M citrate buffer, pH 5.5) and boil for 15 min. Cool the reaction tube, measure the absorbance at 570 nm and calculate the BSH enzyme activity by substituting the formula for the taurine standard curve.

### Molecular docking analysis

To investigate the probable binding of TUDCA to KEAP1 as the potential inhibitor, the automated docking studies were carried out using AutoDock vina 1.1.2 package. The X-ray crystal structure of the human KEAP1 (PDB ID: 6LRZ) was downloaded from RCSB Protein Data Bank. The 3D structure of TUDCA (ZINC ID: 3914813) was downloaded from ZINC. The AutoDockTools 1.5.6 package was employed to generate the docking input files. The molecular docking simulation protein of Keap1 was prepared by removing water molecules and bound ligands. The binding site of the KEAP1 was identified as centre_x: −9.944, centre_y: −38.993, and centre_z: 3.869 with dimensions size_x: 85, size_y: 85, and size_z: 85. The best-scoring (i.e.,with the lowest docking energy) pose as judged by the Vina docking score was chosen and further analyzed using PyMoL 2.4.0 software.

### Statistical analysis

Statistical comparisons among experimental groups were analyzed by one-way ANOVA and Duncan’s multiple-comparison test using the SPSS software (Version 21.0. Armonk, NY: IBM Corp.). *P* values less than 0.05 were considered statistically significant.

## Supporting information

graphic abstract

## Acknowledgements

We gratefully thank Qiaojuan Li and Lingfeng Hou (Wenzhou University) for their assistance in animal experiments. We gratefully thank Alan K Chang (Wenzhou University) for helpful discussion and for revising the language of the manuscript.

## Authors’ contributions

Y.Z., Y.C., J.L, D.H., J.Z. and L.Y performed the research. H.T., Y.Z. and Y.G. designed this study. R.T. and M.W. provided technical assistance. H.T. and Y.Z. analyzed the data and wrote the manuscript. R.T. and Y.G. revised the manuscript. H.T. and Y.G. had primary responsibility for final content. All authors have read and approved the final manuscript.

