## supplementary figures for "Metformin increases tauroursodeoxycholic acid levels to improve insulin resistance in diet-induced obese mice"

**Supplementary figure captions**

**Supplementary Figure 1. Metformin reduces fat accumulation in HFD-fed mice.** (A-H) Mice were fed either normal chow diet (NCD), high fat diet (HFD) or HFD combined with metformin (HFD+Met) throughout 6 weeks. (A) Food intake, (B) organ index, (C) H&E staining of epididymal adipose tissue (400×), (D) number of adipocytes, levels of total cholesterol and triacylglycerol in (F) liver and (G) serum, and (H) serum free fatty acids are shown. Data are expressed as the mean ± SD. *$P < 0.05$, **$P < 0.01$ and ***$P < 0.001$ for HFD *vs* HFD+Met.

**Supplementary Figure 2. Metformin blunts intestinal and liver inflammation.** Determination of inflammatory factors in (A) liver and (B) ileum by *q*PCR, (C) serum LPS by kit. (D) Expression levels of IκB, phosphorylated IκB, P65, and phosphorylated P65 as determined by western blotting and quantitative analysis for the densitometry of p-IκB and p-P65 protein level. Data are expressed as mean ± SD. *$P < 0.05$, **$P < 0.01$ and ***$P < 0.001$ for HFD *vs* HFD+Met.

**Supplementary Figure 3. Metformin remodels fecal microbial structure and communities.** (A) Microbial composition with cluster at the phylum level; (B) relative abundance of gut microbiota at the phylum level; (C) ratio of Bacteroidetes to Firmicutes; (D) function analysis of gut microbiota. (E) Linear discriminant analysis (LDA) score for taxa differing between treatment groups. LDA scores with threshold > 3 indicated a higher relative abundance in the corresponding group than in the other two groups. Data are represented as mean ± SD. $**P < 0.01$ and $***P < 0.001$.

**Supplementary Figure 4. Differential metabolites determined by non-targeted metabolomics.** (A) Number of differential metabolites between each group; (B) the major upregulated differential metabolites in HFD+Met group; (C) the content of TUDCA in the liver/serum; (D) the relative mRNA levels of bile acid synthetase (*Cyp7a1*, *Cyp27a1*, *Baat*) determined by *q*PCR. Data are represented as mean ± SD. $**P < 0.01$ and $***P < 0.001$.

**Supplementary Figure 5. Differential metabolites alleviate insulin resistance.** HepG2 cells were pre-treated with 200 μM PA for 24 h and then incubated with TUDCA (20 μM), taurine (10 mM), or choline hydroxide (10 mM) for 12 h. (A) Glucose uptake and (B) glucose in medium supernatant. (C) Protein levels of p-Akt and Akt were determined by western blotting. Data are expressed as the mean ± SD. $*P < 0.05$, $**P < 0.01$ and $***P < 0.001$.

**Supplementary Figure 6. TUDCA reduces the secretion of pro-inflammatory cytokines.** HepG2 cells were pre-treated with 200 μM PA for 24 h and then incubated with TUDCA (10 or 20 μM) for 12 h. The relative mRNA levels of *Tnf-α*, *Il-1β*, and *Il-6* were determined by *q*PCR. Data are expressed as mean ± SD, $*P < 0.05$.

**Supplementary Figure 7. PA inhibits Nrf2 expression and activation.** PA (200 μM) was incubated with HepG2 cells, and then the protein expression of p-Nrf2 and total Nrf2 was detected by western blotting.

**Supplementary Figure 8. Protein expression levels of Nrf2.** (A) Silencing Nrf2 expression in HepG2 cells by siRNA. Protein levels of Nrf2 were determined by western blotting after 48 hours of transfection with siRNA. (B) Protein levels of Nrf2 in primary hepatocytes from $Nrf2^{-/-}$ mice determined by western blotting.

**Supplementary Figure 9. TUDCA relieves lipid accumulation and intestinal inflammation in *ob/ob* mice.** Total cholesterol and triacylglycerol in (A) serum and (B) liver. (C) The relative mRNA levels of *Tnf-α*, *Il-1β*, and *Il-6* in ileum were determined by *q*PCR. Data are expressed as the mean ± SD. *$P < 0.05$, **$P < 0.01$ and ***$P < 0.001$.

### Supplementary Figure 1

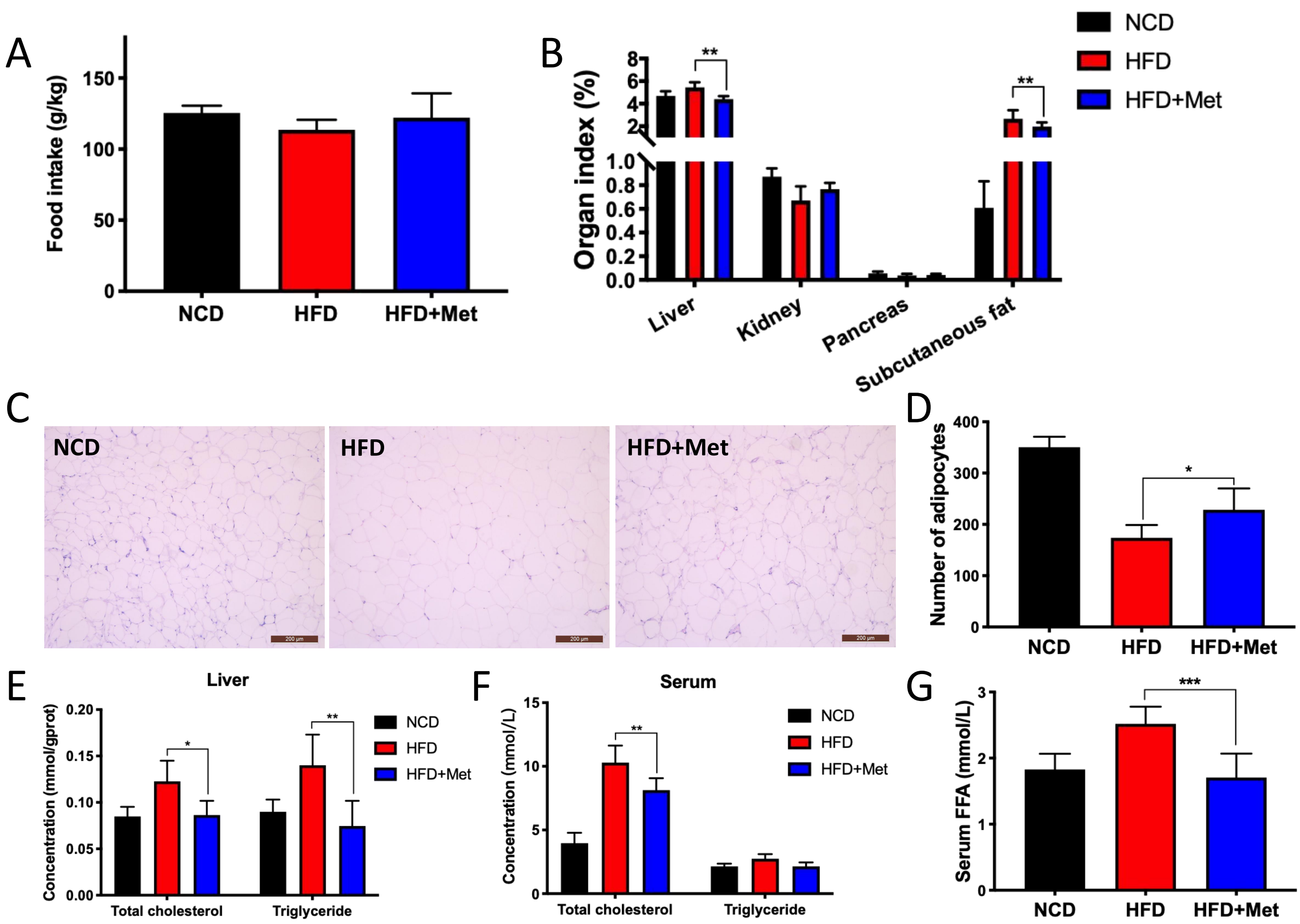

### Supplementary Figure 2

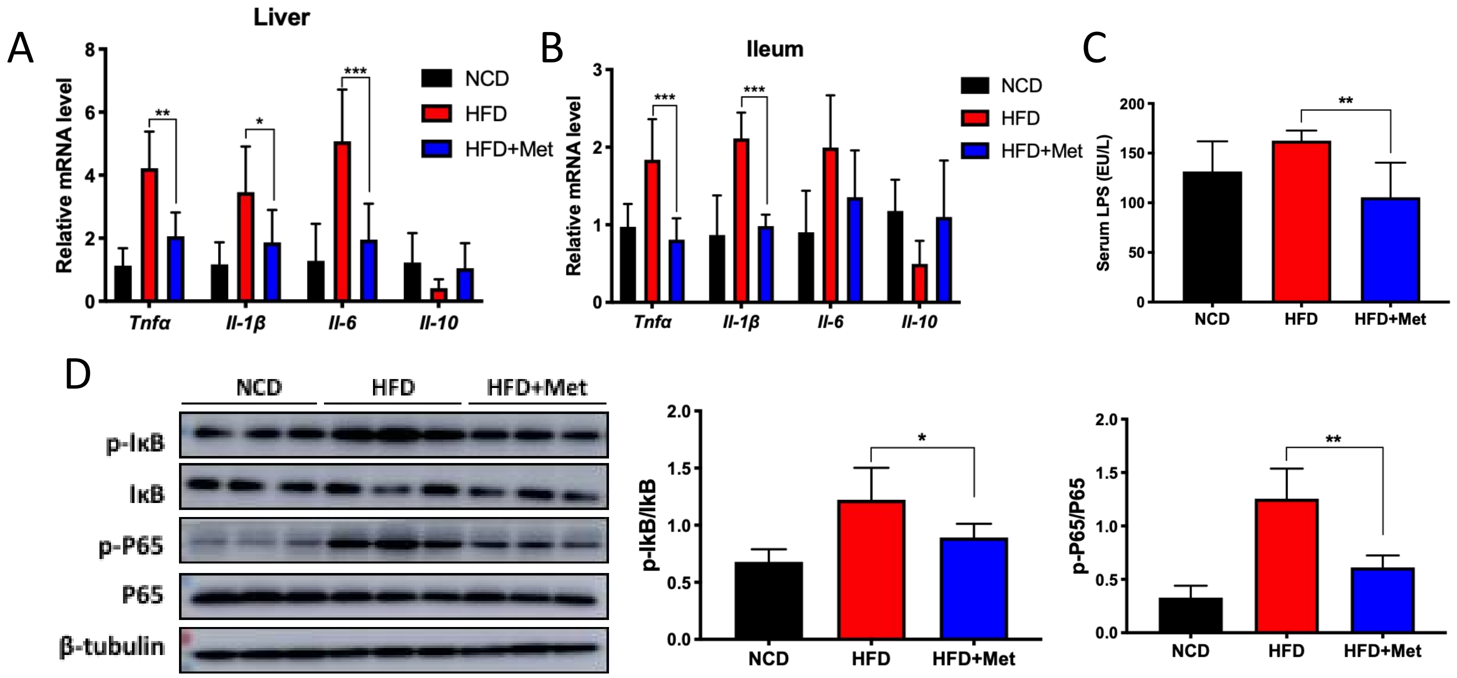

### Supplementary Figure 3

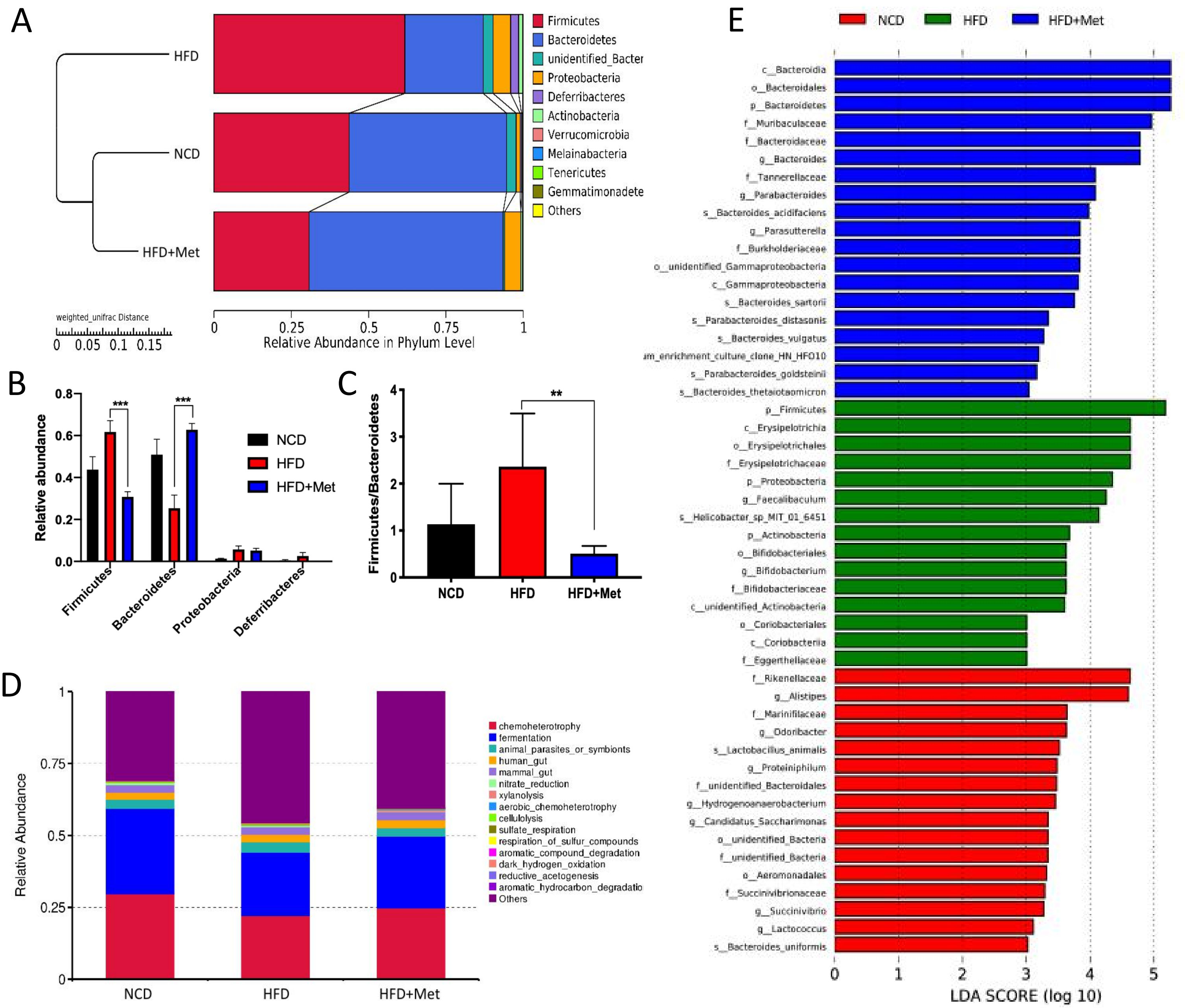

### Supplementary Figure 4

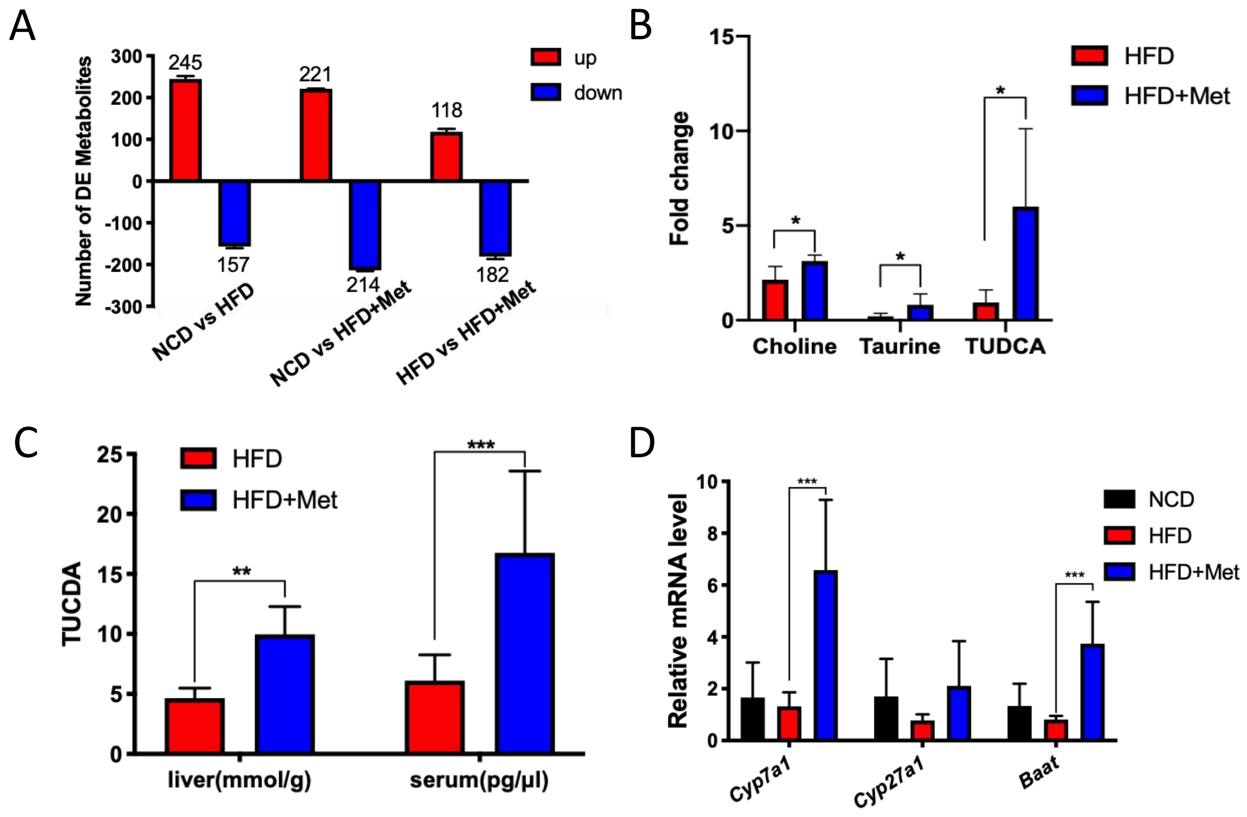

### Supplementary Figure 5

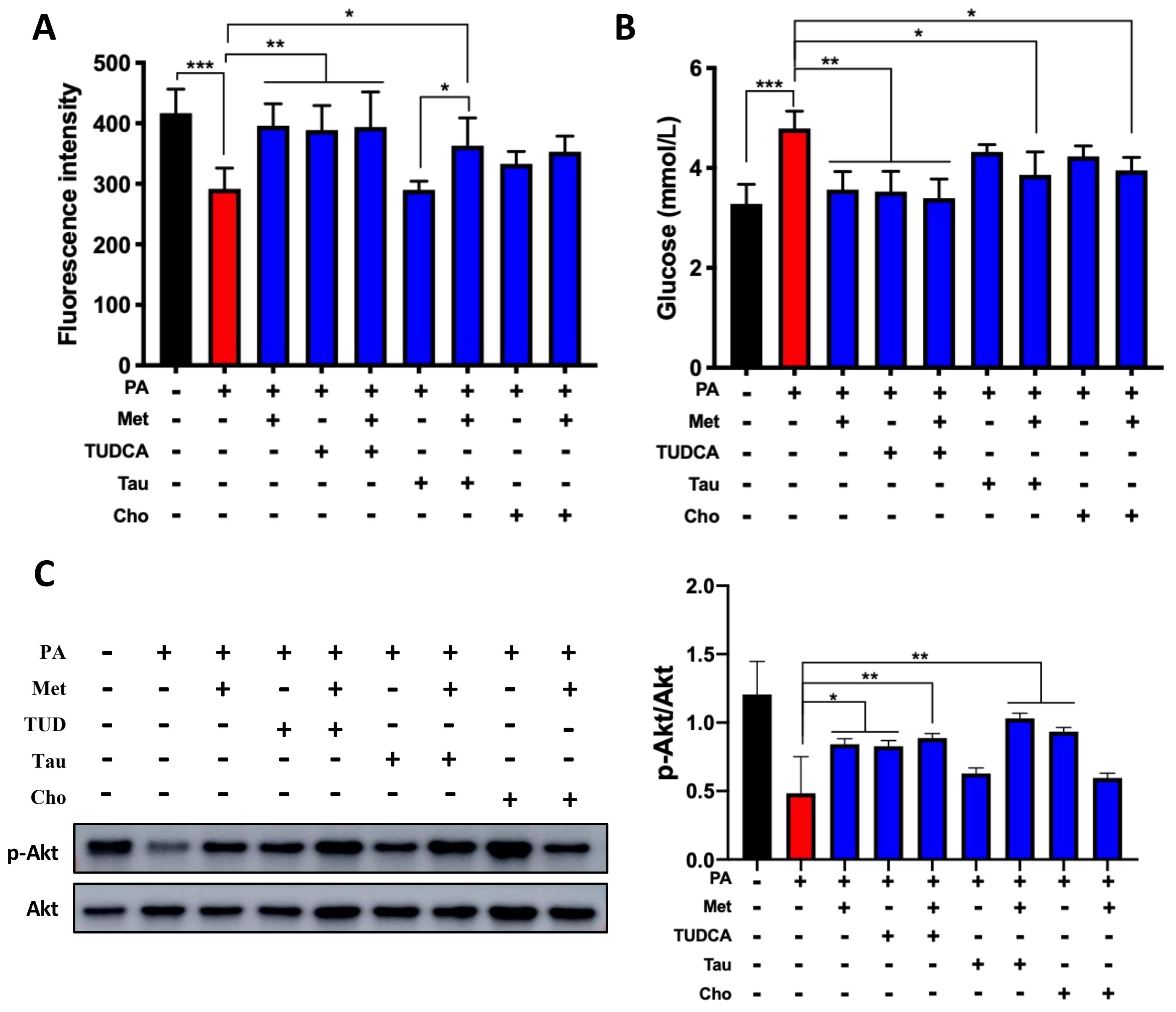

### Supplementary Figure 6

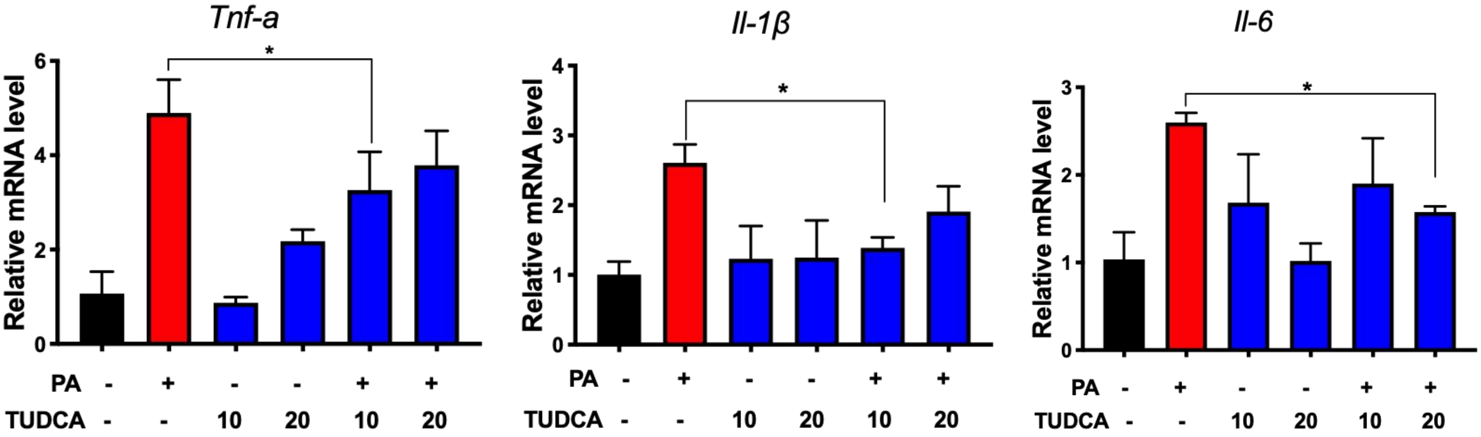

### Supplementary Figure 7

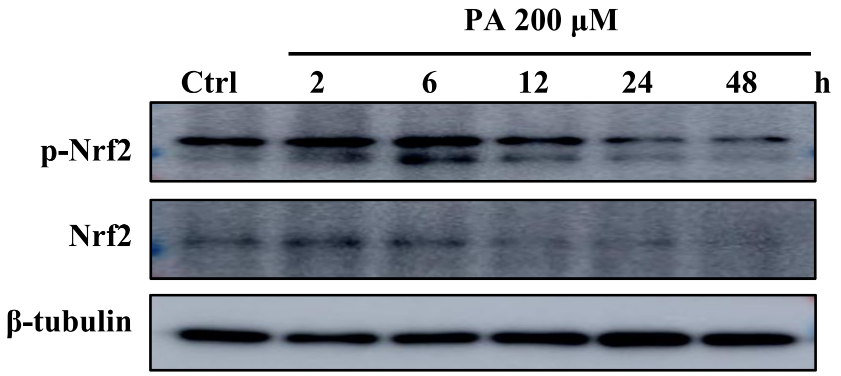

#### Supplementary Figure 8

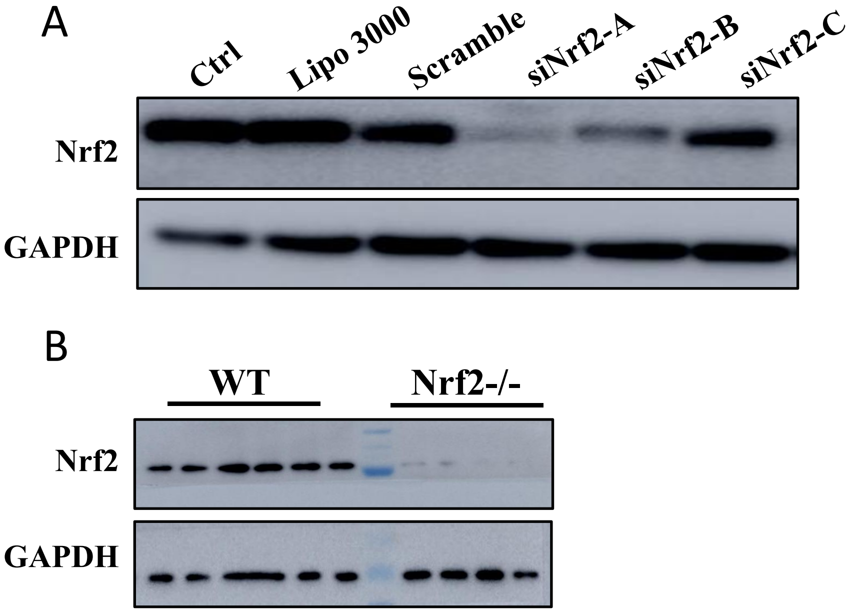

#### Supplementary Figure 9

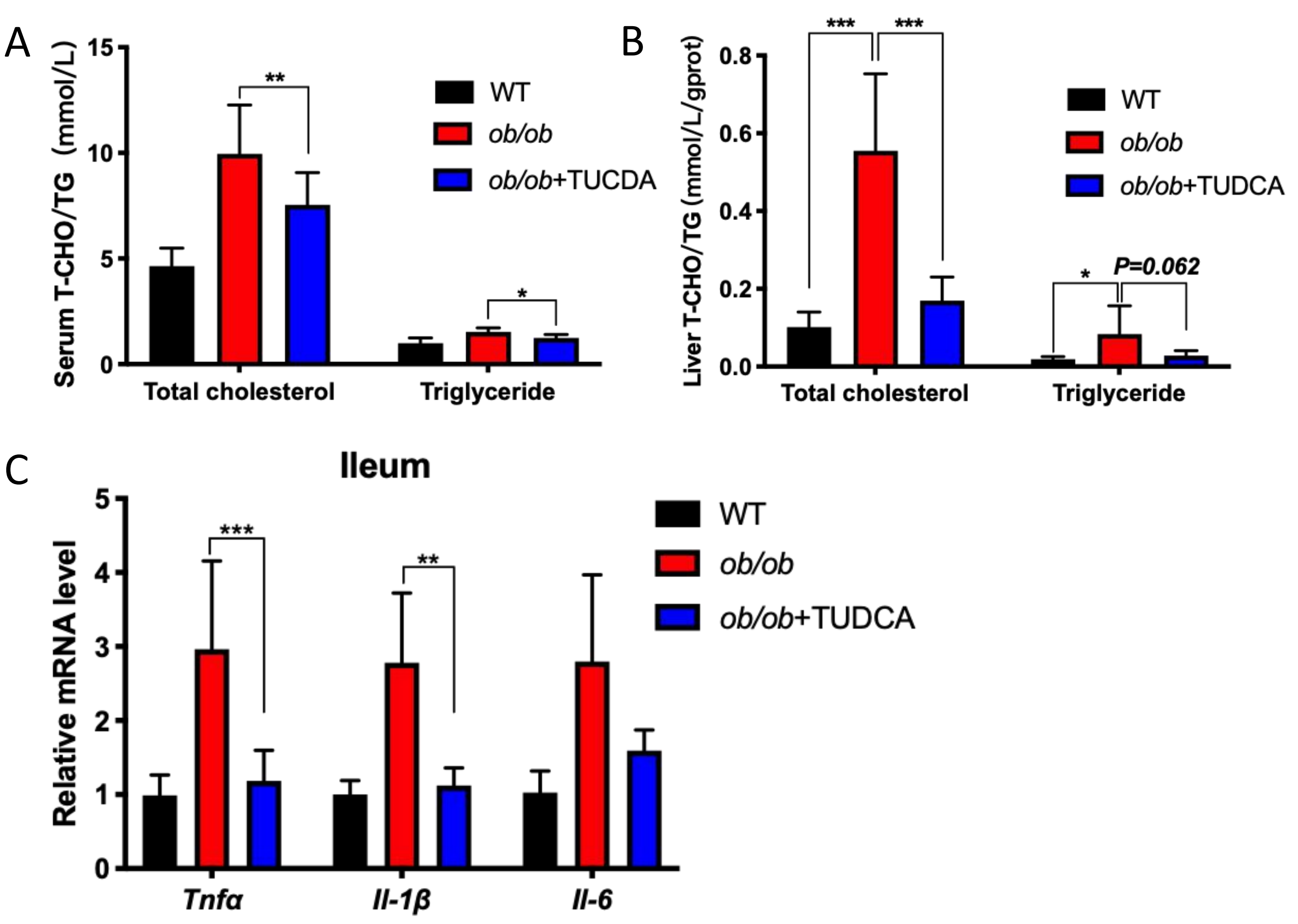
