## supplementary table for "Metformin increases tauroursodeoxycholic acid levels to improve insulin resistance in diet-induced obese mice"

**Supplementary Table. Designed primer sets for qRT-PCR**

| *Gene* | Primer | 5’- 3’ | Size（bp） |
|---|---|---|---|
| Mus *Nrf2* | sense | AAGAATAAAGTCGCCGCCCA | 326 |
| | antisense | GAAAAGGCTCCATCCTCCCG | |
| Homo *Nrf2* | sense | GGTTGGGGTCTTCTGTGG | 247 |
| | antisense | GACATTGAGCAAGTTTGGGAGG | |
| Mus *Tnf-α* | sense | ACCCTCACACTCACAAACCA | 212 |
| | antisense | ATAGCAAATCGGCTGACGGT | |
| Homo *Tnf-α* | sense | CCCTCCAACCCCGTTTTCTC | 157 |
| | antisense | GCCCCTCAAAACCTATTGCC | |
| Mus *Il-1β* | sense | TGCCACCTTTTGACAGTGATG | 220 |
| | antisense | AAGGTCCACGGGAAAGACAC | |
| Homo *Il-1β* | sense | CTCTCAGCAGGTCCGATACC | 198 |
| | antisense | AACATGGCACCTCTGCAACT | |
| Mus *Il-6* | sense | CCCCAATTTCCAATGCTCTCC | 141 |
| | antisense | CGCACTAGGTTTGCCGAGTA | |
| Homo *Il-6* | sense | GGAGTCAGAGGAAACTCAGTT | 210 |
| | antisense | ACTCAGCACTTTGGCATGTCT | |
| Mus *Il-10* | sense | AGGGCACCCAGTCTGAGAACA | 225 |
| | antisense | CGGCCTTGCTCTTGTTTTCAC | |
| Mus *Ho-1* | sense | CACGCATATACCCGCTACCT | 175 |
| | antisense | CCAGAGTGTTCATTCGAGCA | |
| Homo *Ho-1* | sense | CCTTCTTCACCTTCCCCAAC | 124 |
| | antisense | GCCTCTTCTATCACCCTCTG | |
| Mus *Nqo1* | sense | TCACCTGGGCAAGTCCATTC | 241 |
| | antisense | TGCCCTGAGGCTCCTAATCT | |
| Homo *Nqo1* | sense | CAGTTGGGATGGACTTGC | 101 |
| | antisense | CCAGGCAGGATTCTTAATG | |
| Mus *Keap1* | sense | TACACAGCGGGCGGTTACT | 244 |

| | | | |
|---|---|---|---|
| | antisense | TCATAGAGGCACAGGGCGA | |
| Mus *Cyp7a1* | sense | CCGAGTGATGTTTGAAGCCG | 173 |
| | antisense | GGGCTTTATGTGCGGTCTTG | |
| Mus *Cyp27a1* | sense | CCTACATCCATTCGGCTCTG | 145 |
| | antisense | CTTTACTTCTCCCATCCCGG | |
| Mus *Baat* | sense | CTGGAAAGGTGGTATGTGGC | 136 |
| | antisense | ATCAATCCACCAGCACCTCC | |
| Homo *β-actin* | sense | CTCTTCCAGCCTTCCTTCCT | 201 |
| | antisense | TCTTCATTGTGCTGGGTGCC | |
| Mus *β-actin* | sense | CGTGGGCCGCCCTAGGCACCA | 214 |
| | antisense | TTGGCCTTAGGGTTCAGGGGGG | |
| *A. muciniphlia* | sense | CAGCACGTGAAGGTGGGGAC | 128 |
| | antisense | CCTTGCGGTTGGCTTCAGAT | |
| 16s rRNA | sense | GTGACAAACCGGAGGAAGGT | 144 |
| | antisense | ATCCGAACTGAGAACAACTTTATGG | |
