## Supplementary figures and images for "Metformin increases tauroursodeoxycholic acid levels to improve insulin resistance in diet-induced obese mice"

### graphic abstract

#### Graphic abstract

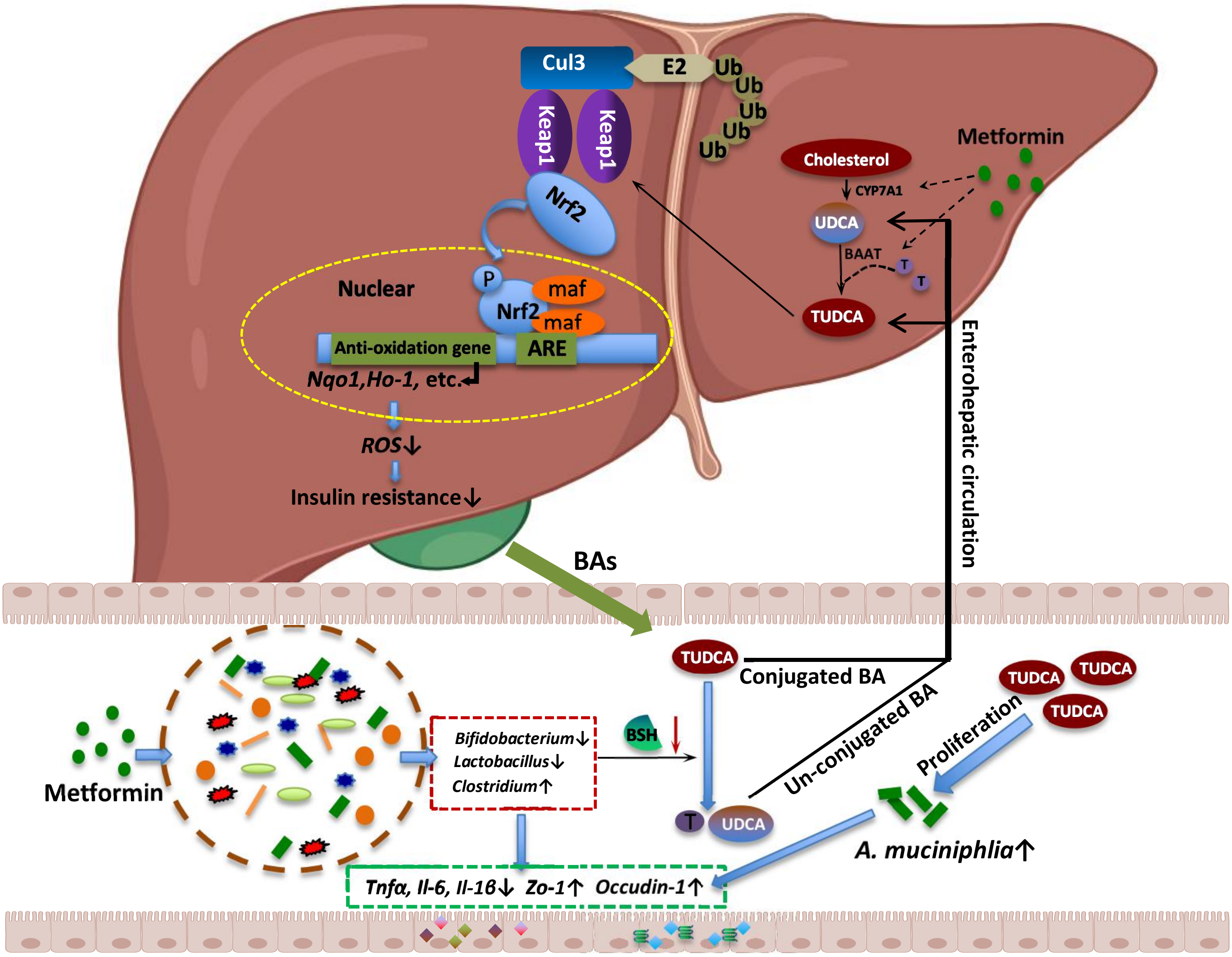
